# Exploring the use of state of nature metrics to screen global business operations for ecological sensitivity and to select priority sites for disclosure

**DOI:** 10.64898/2026.08.25.747065

**Authors:** Elizabeth H. Boakes, Stuart H. M. Butchart, Alena Cierna, Kim Dunn, Julie Dimitrijevic, Frank Hawkins, Owen Jackson, Jodie LeMarquand, Simone Mordue, Richard D. Gregory

## Abstract

Businesses are increasingly encouraged to disclose their nature-related dependencies, impacts, risks and opportunities. A common component of sustainability reporting is screening operational sites for ecologically sensitivity to identify locations for further evaluation and action. However, with 600+ biodiversity metrics available, selecting and interpreting appropriate metrics remains challenging for business. We developed a simple screening framework informed by the Taskforce for Nature-Related Financial Disclosures guidance, grouping eleven widely used global biodiversity metrics into four complementary ‘baskets’, representing different aspects of biodiversity. We created hypothetical but realistic mining, onshore wind energy and agricultural companies, to assess how metric choice, buffer size, scoring approach and sensitivity thresholds influence screening outcomes. Our basket framework consistently identified similar high-priority sites across metric combinations, but site rankings varied with methodological choices. We recommend clearer guidance on metric selection and application, alongside greater transparency from business regarding assumptions, methods and limitations when screening sites for ecological sensitivity

**Short paragraph:** This study explores the use of state of nature metrics to screen global business operations for ecological sensitivity and to inform site prioritization for nature-related financial disclosures.

## 1. Introduction

Economic activities continue to drive the global degradation of biodiversity and ecosystem services, with businesses and their global supply chains contributing to land-use change, climate change, resource extraction, pollution and the spread of invasive alien species, among other threats (Dasgupta, 2021; IPBES, 2019). Yet, these same economic activities driving decline are underpinned by nature. The World Economic Forum estimates that around half of global GDP is highly or moderately dependent on biodiversity (World Economic Forum, 2020) and the recent Intergovernmental Science-Policy Platform on Biodiversity and Ecosystem Services (IPBES) report on Business and Biodiversity states that all businesses both impact and depend on biodiversity (IPBES, 2026). Indeed, biodiversity loss has been identified as one of the most severe global risks facing human society in the next 10 years (World Economic Forum, 2026). Bending the curve of biodiversity loss will require transformative change across governance structures and economic systems (Leclère et al., 2020) and it is increasingly recognised that business has a critical role to play in reversing nature loss (IPBES, 2026). This is reflected in the Kunming-Montreal Global Biodiversity Framework’s (KMGBF) Target 15, (CBD, 2022) which promotes disclosure of biodiversity impacts and dependencies by businesses and financial institutions.

The growing awareness of nature-related risks has led to the development of relevant reporting frameworks and standards for the private sector, for example the Taskforce on Nature-related Financial Disclosures (TNFD) (TNFD, 2023b). The TNFD is a market-led, science-based and government-supported global initiative that is encouraging and enabling organisations to disclose their nature-related dependencies, impacts, risks and opportunities (DIROs). The framework aims for interoperability with other frameworks and reporting standards such as the Science Based Targets Network (SBTN, 2026), the European Sustainability Reporting Standards (ESRS) (European Commission, 2023) and the Global Reporting Initiative (GRI) Standards (GRI, 2023). The aim of these frameworks and standards is to encourage businesses and financial institutions to integrate nature into decision-making, and to enable global financial flows to be directed toward nature-positive outcomes, aligned with the KMGBF. The TNFD framework builds conceptually on the Taskforce for Climate-Related Financial Disclosures (TCFD, 2017). However, reporting on nature is arguably more complex than reporting on climate in part due to a) the complex multiple dimensions of nature, and b) the fact that nature-related DIROs are location-specific. Whereas carbon dioxide emissions are widely accepted as a useful proxy for greenhouse gas emissions, there is no single metric for biodiversity that is universally accepted as adequately reflecting its multiple dimensions, and there is a broad consensus that a suite of metrics is needed (Burgess et al., 2024; CBD, 2022; IPBES, 2019).

The TNFD has developed the LEAP approach – *Locate*, *Evaluate*, *Assess* and *Prepare* – to support organisations to gather and process the information needed to understand their interface with nature throughout their supply chain (TNFD, 2023a). The *Locate* phase facilitates the identification of an organisation’s activities and assets that interact with ecologically sensitive locations and, from these, selecting a manageable number of priority locations for further action. An organisation will then *Evaluate* its dependencies and impacts on nature at these prioritised sites, *Assess* its nature-related risks and opportunities, and *Prepare* to respond and report on material nature-related issues, defined as those that are sufficiently significant to influence assessments and decisions by investors, stakeholders and the organisation itself. A key outcome of the LEAP approach is to influence a business’s capital allocation and land management practices to mitigate the DIROs identified during the disclosure process which often fall across a portfolio of sites across a business’s value chain.

Site-based operations in the extractive, renewable energy and agricultural sectors can impact biodiversity both directly (e.g. via pollution, collisions with turbines and land use change) and indirectly (e.g. via increased access to previously intact forest areas and increased in-migration and settlement) (Narain et al., 2026). It is important that organisations capture the full range of these impacts by applying a distance buffer around a site to represent the organisation’s ‘Area of Influence’ (AoI) (Laurance et al., 2015; TNFD, 2023a). Once a buffer has been applied, organisations then need to choose appropriate nature-based metrics and assess whether their values exceed thresholds that would identify sites as ‘ecologically sensitive’. Finally, they need to interpret the results to decide which of their sites are of highest priority for nature and should be examined in more detail at the *Evaluate* phase. The decisions required at the *Locate* phase are not straightforward and the spatial heterogeneity, scale-dependence and general uncertainty surrounding the relationships between biodiversity and ecosystem services (Mace et al., 2018; Nicholson et al., 2021) mean there is no one-size-fits-all solution (Narain et al., 2026; Newbold et al., 2016).

A variety of resources exist to assist businesses. Arguably the most scientifically credible of these is the Integrated Biodiversity Assessment Tool (IBAT) (IBAT, 2026) which offers a TNFD disclosure service for organisations that includes guidelines around buffer sizes for different sectors’ operations, defined thresholds for proximity to Protected Areas (PAs), proximity to Key Biodiversity Areas (KBAs) and the Species Threat Abatement and Restoration (STAR) metric (a measure of the potential contribution that specific conservation actions at particular locations can make toward reducing the global extinction risk of species (Mair et al., 2021)), and a scoring system to allow metrics to be combined to produce a total score for ranking areas’ biodiversity importance (IBAT, 2024). However, some organisations use other platforms and tools without such datasets and services, and organisations also need to interpret metrics related to ecosystem integrity, water risk etc. The LEAP guidance stipulates organisations should report their decision-making process at the Locate phase (TNFD, 2023a) but, to date, few disclosures contain this information. This lack of transparency means it is impossible to gauge how organisations have been making these decisions or whether they are appropriate.

One practical question that remains unanswered is the degree to which factors such as metric choices, buffer size, sensitivity threshold, and metric scoring system, determine sites’ ecological sensitivity rankings. Here we address this gap by investigating how much influence these considerations have on site rankings and therefore the decisions made by businesses to protect and recover nature. We created three hypothetical companies’ operational site portfolios in the mining, onshore wind energy and agriculture sectors and explored how; a) metric choice, b) buffer size, c) metric sensitivity thresholds, and d) metric scoring systems affected the ranking of sites with respect to ‘areas important for biodiversity’ and ‘areas of high ecosystem integrity’, the two *Locate* criteria that directly relate to the TNFD’s ‘State of Nature’ disclosure metrics.

## 2. Methods

### 2.1 Creating direct operations sites

We created operational site portfolios for three imaginary companies to explore the decisions businesses take during the TNFD’s L4 process of *Locate* (or similar location screening processes for GRI, SBTN etc.) Each company represented a sector with a particularly high impact on nature (Finance for Biodiversity Foundation, 2023) - Mining, Energy (represented by onshore wind energy) and Agriculture. (For simplicity we considered direct operation sites only although the TNFD requires organisations to assess their entire supply chain.) Sites were selected from global-scale datasets of mining areas (Maus et al., 2020), wind farms (Dunnett et al., 2020) and croplands (Tang et al., 2024) to ensure they were in realistic locations. To make the companies as globally representative as possible, we spread sites across the following countries, chosen to represent a range of geographies and biomes: Australia, Brazil, Canada, Democratic Republic of Congo (DRC), the United Kingdom (UK), Indonesia and South Africa. We aimed for 10 sites in each country, but our onshore wind portfolio differed slightly with no sites in DRC and only 5 sites in Indonesia due to a lack of wind farms in these countries. Within each country, we ensured sites represented a range of sizes and that overlap between each site’s buffer or ‘Area of Influence’ (AoI), was avoided if possible or otherwise minimised.

We discarded very small sites from the global datasets of mining and renewable energy sites (< 0.5 km^2^ (Mining), <3 turbines (Onshore wind energy)) and ordered the remainder of the sites by size within each country. We then divided sites into size deciles and picked one site at random from each size decile to ensure we covered a representative range of site sizes. Using the ‘terra’ package in R (Hijmans et al., 2026), we applied buffers to each site (following IBAT buffer size guidelines of 50 km for Mining and 10 km for Onshore wind energy (IBAT, 2024)) and replaced any sites with overlapping buffers. It was not possible to pick 10 non-overlapping buffered Mining sites in the UK due to its smaller size and so UK sites were hand-selected to minimise buffer overlap. Neither was it possible to avoid overlapping buffered Onshore wind energy sites in Indonesia as there were only 5 sites in total, 4 of which overlapped.

For Agriculture, we selected sites associated with each country’s largest export crop: wheat - Australia, soy - Brazil, canola (rapeseed) - Canada, robusta coffee - DRC, barley - UK, oil palm - Indonesia and citrus – South Africa (FAO, 2026). We used the ‘Crop area’ layer of the CROPGRIDS dataset (Tang et al., 2024) which gives the actual land area cultivated for a specific crop, regardless of whether it is harvested once or multiple times a year. The crop data were in raster format at a resolution of ∼5.5 km^2^ so we sampled the grid cells at random to make realistic field sizes based on Lesiv et al.’s (2019) estimates of the global distribution of field sizes (very small < 0.64 ha; small 0.64-2.56 ha; medium 2.56-16 ha; large 16-100 ha; very large >100 ha). In each country, we selected 10 cropland cells at random, took the centroid of the cell, and then drew a circle around it to create an area equivalent to the cropland area cultivated within that cell. In each country, we made three fields in the small range, three in the medium, two in the large and two in the very large, to ensure sites represented a range of sizes. We did not make any ‘very small’ fields as these would be less likely to be relevant for global companies. We assumed that agriculture in DRC was low-intensity (given this is typically the case) but that sites in all other countries were high-intensity and, following IBAT’s guidelines (IBAT, 2024), applied a 5 km buffer to each site in DRC, and a 10 km buffer to all other sites, replacing any sites with overlapping buffers.

### 2.2 Metrics

Metrics used at *Locate* should ideally be comparable across a wide range of geographies, accessible, timely, location-explicit and useful in different business contexts and across sectors. Table 4 of the TNFD’s LEAP guidance (TNFD, 2023a) lists recommended metrics and reference datasets that can be used to identify sensitive locations for each of five different sensitivity criteria. Additional options are given in Annex 2 of the LEAP guidance (TNFD, 2023a), the short list of the Nature Positive Initiative’s ‘Spreadsheet: Terrestrial Metrics for Consultation’ (Nature Positive Initiative, 2026) and by Burgess et al. (2024). We chose to explore metrics from these three sources that have terrestrial global coverage, are fully developed and tested (some metrics listed in Annex 2 are still in development), are available at a resolution of 10km^2^ or finer, are suitable for the *Locate* stage, and relate to the *Locate* criteria of areas important for biodiversity or areas important for ecosystem integrity (Table 1). Further information on the metrics is given in the Supplementary Information (Appendix A) where we summarise each metric’s key features, limitations and underlying data. Appendix B gives information on metrics’ attributes (e.g. units, resolution, year of most recent release) and qualities (e.g. taxonomic and geographic representativeness, conservation action relevance). We did not examine the TNFD’s criteria of ‘areas of rapid decline in ecosystem integrity’ since ‘rapid’ is as yet undefined with respect to both time period and extent of decline.

**Table 1.**
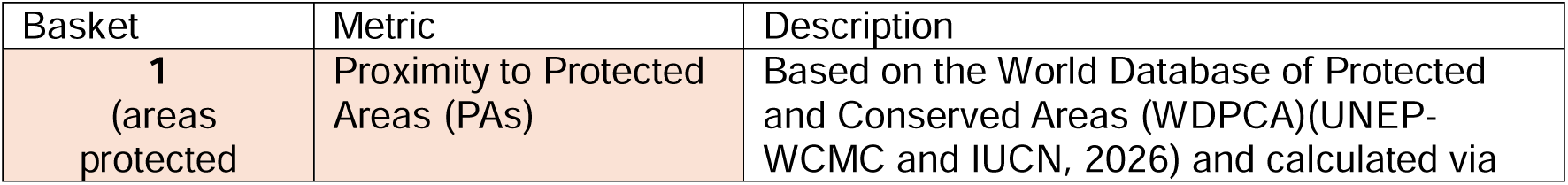
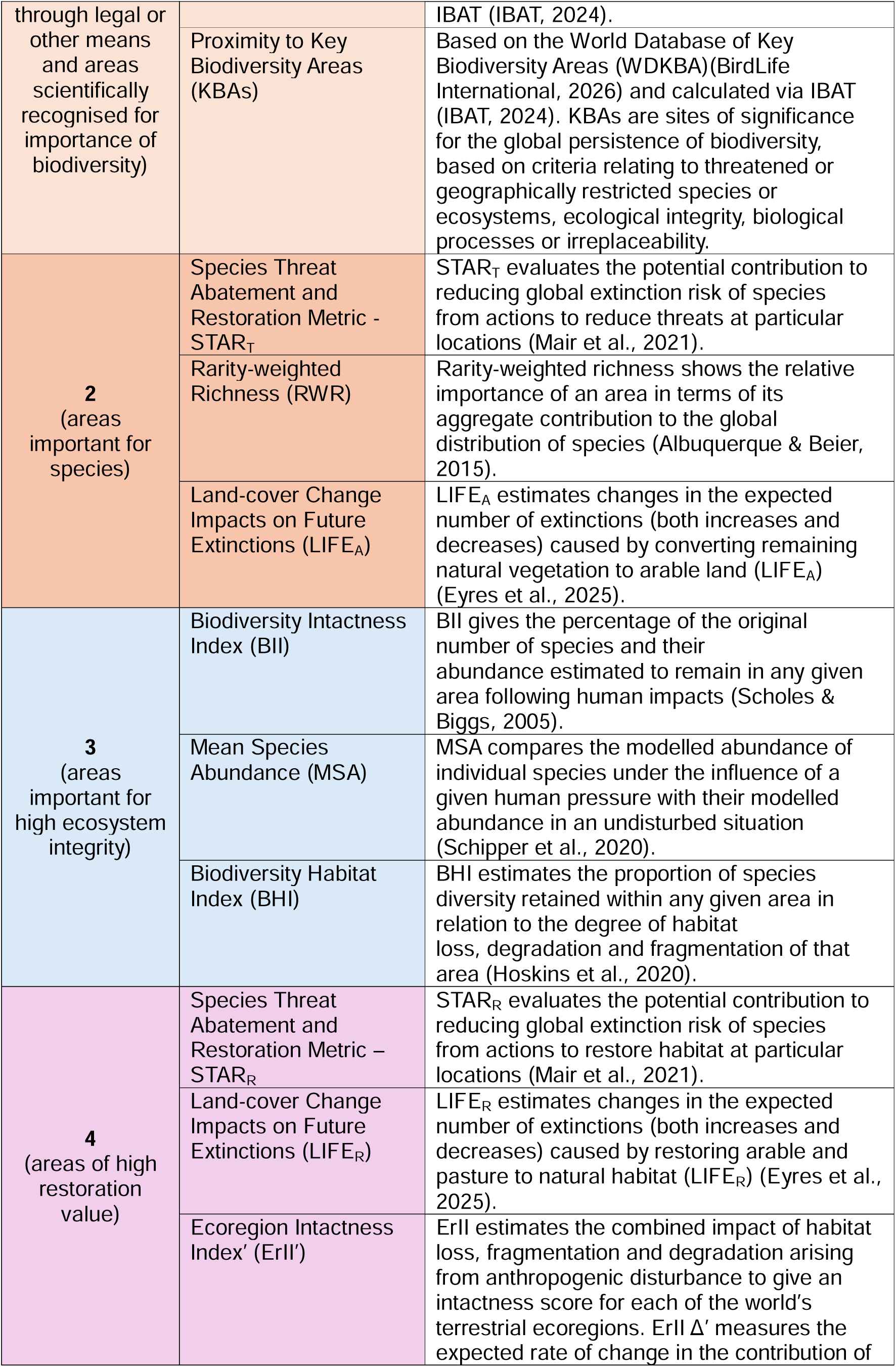
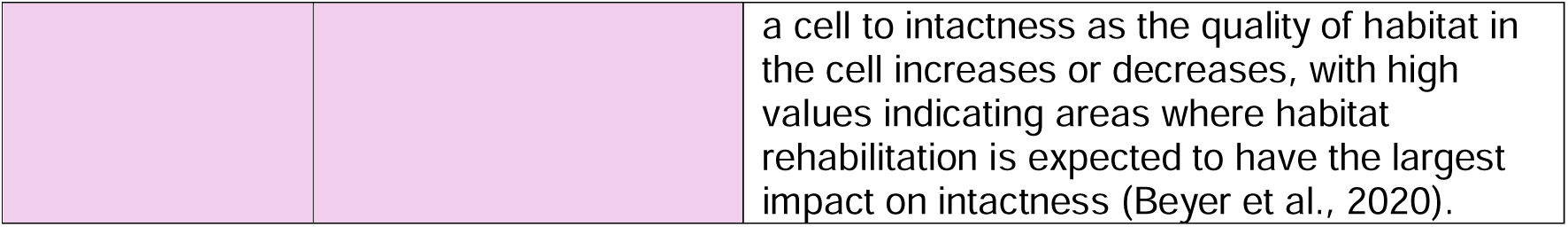
A description of the metrics used in the analysis and their allocation into ‘Baskets’. Shading differentiates metrics used to measure areas of biodiversity importance (light and medium brown corresponding to Baskets 1 and 2, respectively), areas of high ecosystem integrity (blue - Basket 3) and areas of high restoration value (pink – Basket 4).

### 2.3 Metric Basket Protocol

Many of the metrics being promoted for private sector biodiversity disclosures rely on the same or similar underlying data (Constantino-Panopio et al., 2025) and are highly correlated with each other both spatially and temporally. Hence, we suggest a structured framework to help simplify metric choice for businesses at the Locate stage whilst ensuring key dimensions of biodiversity are measured with minimal redundancy (i.e. where multiple metrics reflect largely the same dimension of biodiversity). We call these groups of metrics ‘baskets’ (see Table 1 and Figure 2). Our structured framework is based on the TNFD’s criteria for sensitive location identification and aims to create complementary combination of metrics with low redundancy.

**Figure 1.**
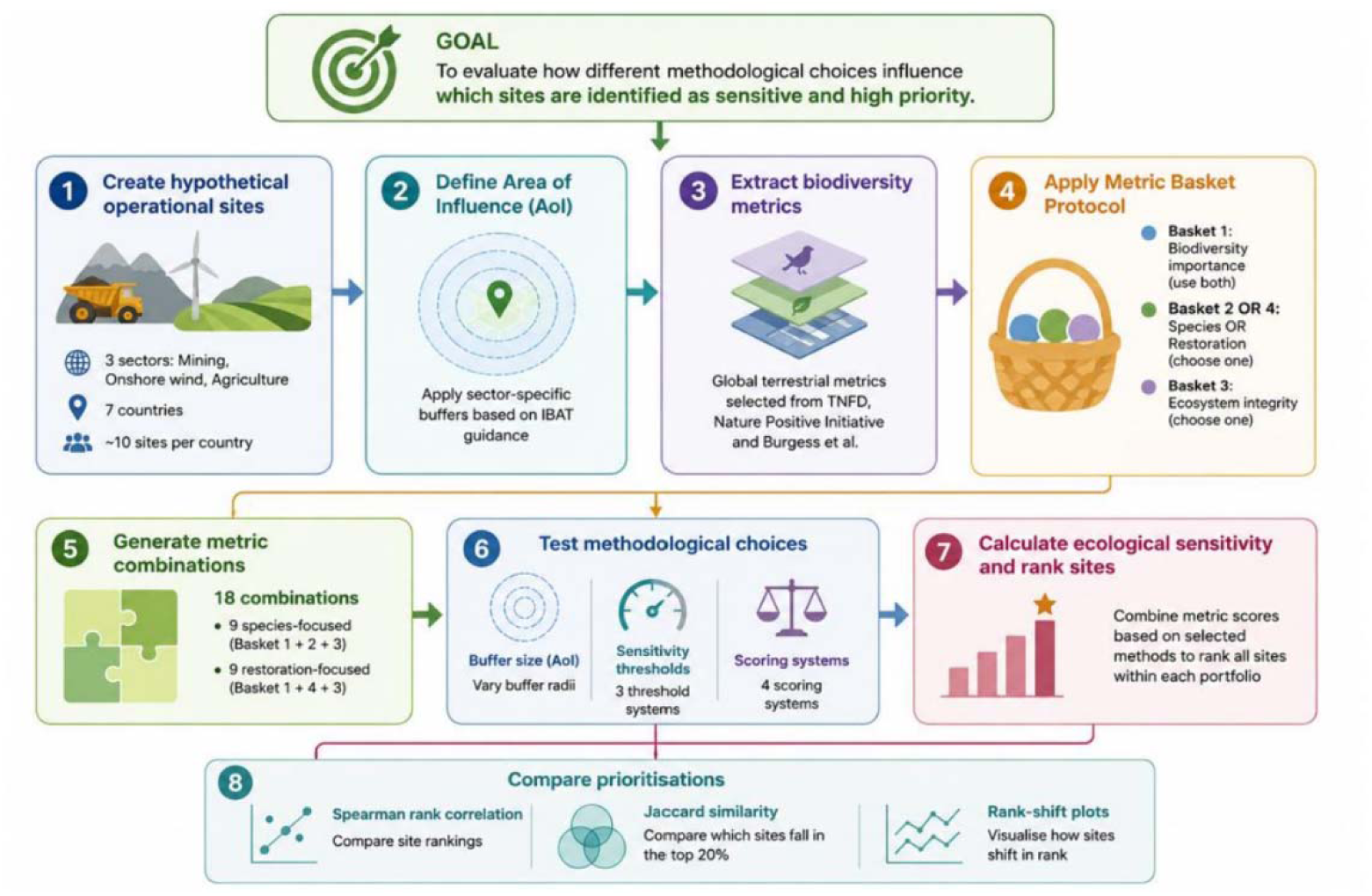
A visual overview of the Methods.

**Figure 2.**
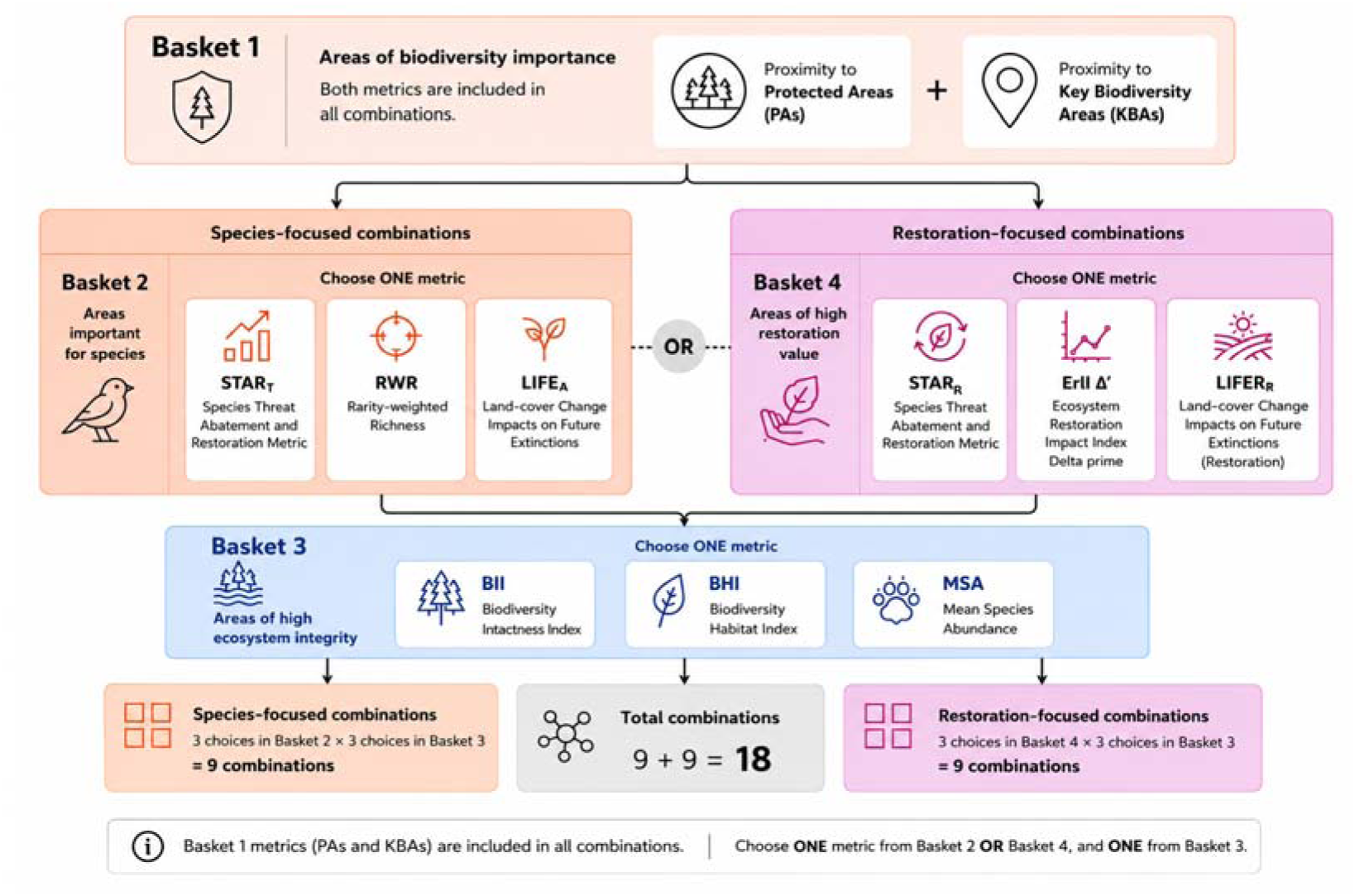
A visual representation of the Basket Framework.

#### User must pick **both** metrics from Basket 1: Proximity to PAs and proximity to KBAs

These two metrics reflect distance from delineated sites of conservation or biodiversity importance. Protected areas are sites with statutory, regulatory status and may not all be of high biodiversity significance. KBAs are sites of international significance for biodiversity and may not all be officially protected. Hence these two metrics are complementary and should not be considered interchangeable. We therefore suggest that users assess both metrics.

#### User must pick **one** metric from Basket 2: STAR_T_, RWR, LIFE_A_

These three metrics identify locations that are particularly important for global biodiversity as they support species at risk of extinction and often contain a substantial proportion of those species’ worldwide ranges. All three metrics are derived from species distribution data from the IUCN Red list. As they measure very similar aspects of biodiversity importance, using more than one of these would be largely redundant for the purposes of the *Locate* step. Nevertheless, users should consider which metric is most appropriate for their needs with STAR_T_ indicating priority areas for implementing conservation actions to abate threats as well as the individual threats to focus on (Mair et al., 2021) and LIFE_A_ indicating priority areas for habitat protection with respect to conversion to agricultural land (Eyres et al., 2025). RWR does not contain information on either individual species or threats (IUCN, 2025).

#### User must pick **one** metric from Basket 3: BII, BHI, MSA

These metrics are all based on models of how species’ populations respond to anthropogenic threats and are used as proxies of ecosystem integrity with respect to species composition, which is a form of measuring ecosystem condition. Using more than one creates duplication and is unnecessary for the purposes of *Locate.* However, as for Basket 2, when picking a metric users should consider their key features and limitations (see Supplementary Information, Appendix A). BII and BHI and modelled on a much wider range of taxa and geographies than MSA’s GLOBIO4, for example.

#### If screening areas for ecosystem restoration value, as an alternative to Basket 2, users may pick **one** metric from Basket 4: STAR_R_, LIFE_R_, ErII Δ’

Identifying areas for restoration opportunities is not included within the TNFD’s *Locate* process but there are scenarios where businesses might wish to choose sites for restoration activities (e.g. local community projects or mine closure planning) and screening localities for restoration value would be a useful and straightforward initial step for business when researching conservation opportunities within LEAP. Basket 4 metrics relate to identifying opportunities for restoring habitat as opposed to Basket 2 metrics which focus on the negative impact of business activities.

STAR_R_ quantifies the reduction in global extinction risk that could be achieved from restoring habitat (Mair et al., 2021). LIFE_R_ estimates the change in extinction probability resulting from restoring agricultural land and so would not be appropriate to evaluate land used for other purposes (Eyres et al., 2025). ErII Δ’ measures the expected rate of change in the contribution of a spatial unit (grid cell) to intactness as the quality of habitat in the grid cell changes (Beyer et al., 2020). STAR_R_ and LIFE_R_ are both derived from data on species’ distributions from the IUCN Red List (IUCN, 2025), whereas ErII Δ’ is derived from the Human Footprint Index (Venter et al., 2016) and data on ecoregions (Dinerstein et al., 2017). Using more than one of these metrics creates duplication and is unnecessary for the purposes of an initial screening of sites.

### 2.4 Metric combinations

Following the Metric Basket Protocol detailed above, we explored 18 different combinations of metrics. 9 combinations assessed sites using metrics from Basket 1 (areas protected through legal or other means and areas scientifically recognised for importance for biodiversity), Basket 2 (areas important for species), and Basket 3 (areas of high ecosystem integrity). The remaining 9 combinations used metrics from Basket 1, Basket 4 (areas important for restoration) and Basket 3. The metric combinations are listed in Table 2.

**Table 2.** The 18 metric combinations used in the analysis.

| Combination | Basket 1 | Basket 2 | Basket 3 | Basket 4 |
| --- | --- | --- | --- | --- |
| A | Proximity to PAs,<br>Proximity to KBAs | STAR<sub>T</sub> | BII |  |
| B | Proximity to PAs,<br>Proximity to KBAs | STAR<sub>T</sub> | BHI |  |
| C | Proximity to PAs,<br>Proximity to KBAs | STAR<sub>T</sub> | MSA |  |
| D | Proximity to PAs,<br>Proximity to KBAs | RWR | BII |  |
| E | Proximity to PAs,<br>Proximity to KBAs | RWR | BHI |  |
| F | Proximity to PAs,<br>Proximity to KBAs | RWR | MSA |  |
| G | Proximity to PAs,<br>Proximity to KBAs | LIFE<sub>A</sub> | BII |  |
| H | Proximity to PAs,<br>Proximity to KBAs | LIFE<sub>A</sub> | BHI |  |
| J | Proximity to PAs,<br>Proximity to KBAs | LIFE<sub>A</sub> | MSA |  |
| K | Proximity to PAs,<br>Proximity to KBAs |  | BII | STAR<sub>R</sub> |
| L | Proximity to PAs,<br>Proximity to KBAs |  | BHI | STAR<sub>R</sub> |
| M | Proximity to PAs,<br>Proximity to KBAs |  | MSA | STAR<sub>R</sub> |
| N | Proximity to PAs,<br>Proximity to KBAs | | BII | ErII $\Delta'$ |
| P | Proximity to PAs,<br>Proximity to KBAs | | BHI | ErII $\Delta'$ |
| Q | Proximity to PAs,<br>Proximity to KBAs | | MSA | ErII $\Delta'$ |
| R | Proximity to PAs,<br>Proximity to KBAs |  | BII | LIFE<sub>R</sub> |
| S | Proximity to PAs,<br>Proximity to KBAs |  | BHI | LIFE<sub>R</sub> |
| T | Proximity to PAs,<br>Proximity to KBAs |  | MSA | LIFE<sub>R</sub> |

### 2.5 Area of Influence/buffer size

For all metrics, we explored the effect of buffer size on the number of sites identified as ecologically sensitive by varying buffer size at 1km intervals from 0-70 km (for Mining sites) and 0-20 km (for Onshore wind energy and Agriculture sites). We examined how buffer size affects the ranking and prioritisation of sites with respect to ecological sensitivity, comparing IBAT’s sectoral buffer size recommendations (IBAT, 2024) with buffer radii 50% and 80% smaller. Concentric buffers were added around site boundaries using the terra package in R (Hijmans et al., 2026).

### 2.6 Sensitivity Thresholds

We used three different systems to assign sensitivity thresholds. Two were objective and based on percentiles of the global datasets. The third was based, where possible, on published sensitivity thresholds associated with each metric, or on thresholds that we chose based on methods mirroring those published (see Table 3).

**Table 3.** The sensitivity thresholds used for each metric.

| Metric | Sector | Lower sensitivity | Medium sensitivity | High sensitivity | Threshold rationale |
| --- | --- | --- | --- | --- | --- |
| Proximity to PAs | Mining | 15-50 km | 5 - < 15 km | < 5 km | IBAT guidelines (IBAT, 2024) |
|  | Onshore wind energy/<br>Agriculture | 3-10 km | 1 - < 3 km | < 1 km |  |
| Proximity to KBAs | Mining | 15-50 km | 5 - < 15 km | < 5 km |  |
|  | Onshore wind energy/<br>Agriculture | 3-10 km | 1 - < 3 km | < 1 km |  |
| STAR<sub>T</sub> | All | $7.95 * 10^{-3} - < 3.61 * 10^{-2}$ | $3.61 * 10^{-2} - < 0.151$ | $\geq 0.151$ | Upper median |
| | | $3.61 * 10^{-2} - < 0.151$ | $0.151 - < 0.438$ | $\geq 0.438$ | Upper quartile |
| | | $0.01 - < 0.05$ | $0.05 - < 0.15$ | $\geq 0.15$ | IBAT guidelines |
| RWR | All | $5.53 * 10^{-2} - < 0.171$ | $0.171 - < 0.370$ | $\geq 0.370$ | Upper median |
| | | $0.171 - < 0.370$ | $0.370 - < 0.662$ | $\geq 0.662$ | Upper quartile |
| | | $0.266 - < 0.370$ | $0.370 - < 0.662$ | $\geq 0.662$ | Sensitivity threshold set at 85<sup>th</sup> percentile as this is where Albuquerque & Gregory (2017) identified the cumulative curve of values flattening. We set medium and high thresholds at 90<sup>th</sup> and 95<sup>th</sup> percentiles |
| LIFE<sub>A</sub> | All<br>(most likely only relevant for Agricultural sector) | $5.83 * 10^{-6} - < 2.54 * 10^{-5}$ | $2.54 * 10^{-5} - < 7.44 * 10^{-4}$ | $\geq 7.44 * 10^{-4}$ | Upper median |
| | | $2.54 * 10^{-5} - < 7.44 * 10^{-5}$ | $7.44 * 10^{-5} - < 1.43 * 10^{-4}$ | $\geq 1.43 * 10^{-4}$ | Upper quartile |
| | | $1.94 * 10^{-5} - < 3.54 * 10^{-5}$ | $3.54 * 10^{-5} - < 7.44 * 10^{-5}$ | $\geq 7.44 * 10^{-5}$ | Sensitivity threshold set at 70<sup>th</sup> percentile as, following Albuquerque & Gregory's (2017) methodology, this is approximately where the cumulative curve starts to flatten. We set medium and high thresholds at 80<sup>th</sup> and 90<sup>th</sup> percentiles |
| BII | All | 57.9 - < 79.9 | 79.9 - < 96.2 | $\geq 96.2$ | Upper median |
|  |  | 79.9 - < 96.2 | 96.2 - < 98.7 | >= 98.7 | Upper quartile |
|  |  | 80 - < 85 | 85 - <90 | >=90 | Newbold et al. (2016) suggest the planetary boundary for BII should be at 90 with a lower value of 80 to cover uncertainty. We took these to be the high and low boundaries and set the medium threshold to be equally spaced between low and high. |
| BHI | All | 0.726 - < 0.796 | 0.796 - < 0.861 | >= 0.861 | Upper median |
|  |  | 0.796 - < 0.861 | 0.861 - < 0.884 | >= 0.884 | Upper quartile |
|  |  | 80 - < 85 | 85 - <90 | >=90 | Since BHI is based on BII values we used the same thresholds as for BII |
| MSA | All | 0.546 - < 0.831 | 0.831 - < 0.872 | >= 0.872 | Upper median |
|  |  | 0.831 - < 0.872 | 0.872 - < 0.880 | >= 0.880 | Upper quartile |
|  |  | 0.720 - < 0.810 | 0.810 - < 0.900 | >= 0.900 | Lucas & Wilting (2018) calculate a planetary boundary for MSA based on the MSA value that corresponds to a BII value of 80. We chose medium and high values to be equally spaced between low and 100. |
| STAR<sub>R</sub> | All | 0.003 -< 0.02 | 0.02 -<0.05 | >=0.05 | IBAT guidelines |
| LIFE<sub>R</sub> | All | $-5.85 * 10^{-5} - < 1.7 * 10^{-4}$ | $-1.7 * 10^{-4} - < -0.32 * 10^{-4}$ | $>= -0.32 * 10^{-4}$ | Upper quartile |
| ErII Δ' | All | 2228 - < 2546 | 2546 - < 2808 | >=2808 | Upper quartile |

#### Upper median

Thresholds were based on 50^th^, 75^th^, 90^th^ percentiles of global values (see Table 3 for values).

#### Upper quartile

Thresholds were based on 75^th^, 90^th^, 95^th^ percentiles of global values (see Table 3 for values).

#### Published/derived thresholds

Ideally, ecosystem sensitivity thresholds would correspond to the levels of disturbance at which ecosystem function begins to break down. For example, the authors of the Forest Landscape Integrity Index (not used in this analysis) categorised scores into low, medium and high integrity by benchmarking against reference locations of forests worldwide (Grantham et al., 2020). However, there is much uncertainty around the resilience of ecosystems and it is likely that resilience will not be uniform across the globe (Mace et al., 2018; Newbold et al., 2016; Nicholson et al., 2021). While the authors of some of the metrics we used have suggested (heavily caveated) values which would correspond with high integrity ecosystem, in other instances, sensitivity thresholds have been derived by examining a curve of ranked values (Albuquerque & Gregory, 2017) or by benchmarking values to a different metric for which a threshold has previously been proposed (Lucas & Wilting, 2018). Other metrics have sensitivity thresholds recommended by IBAT (IBAT, 2024). For those that did not fall into any of these categories, we approximated values ourselves. Sensitivity thresholds and the logic behind each are given in Table 3. We emphasise that we are not endorsing these values nor offering an opinion on their validity, we are simply exploring their use in disclosures.

### 2.7 Scoring Systems

There are a variety of ways to combine metrics to calculate total ecological sensitivity scores as an aid to site prioritisation, each with strengths and weaknesses. We examine four:

**Simple additive** – a linear scoring system was used to score no (0), low (1), medium (2) and high (3) sensitivity with scores then added across metrics. The advantage of the simple additive system is that it is straightforward and intuitive for business to understand.

**Weighted additive** - a linear scoring system was used to score no (0), low (1), medium (2) and high (3) sensitivity. Mean scores were calculated for biodiversity importance metrics and, separately, for ecosystem integrity metrics. We used an equal weighting system to calculate the final score. Weighted additive score = (mean Biodiversity Importance score * 0.5) + (mean Ecosystem integrity score * 0.5).

The advantage of this system is that it gives equal weighting to each of the two major dimensions in our Locate criterion. It could be adapted to incorporate other dimensions of ecological sensitivity and to weight particular criteria.

**Multiplicative** – a linear scoring system was used to score no (1), low (2), medium (3) and high (4) sensitivity. (Note that a score of zero should not be assigned to any metric as it will lead to a combined score of zero.) Mean scores were then calculated for biodiversity importance metrics and, separately, for ecosystem integrity metrics. These two scores were then multiplied to give the final score. Multiplicative score = mean Biodiversity Importance score * mean Ecosystem Integrity score.

The advantage of this system is that sites that are sensitive for two criteria (i.e. biodiversity importance **and** ecosystem integrity) score higher than those sensitive for only one.

**Trigger** – a polynomial-based scoring system was used to score no (0), low (1), medium (2*^n^*) and high (3*^n^*) sensitivity, where *n* is the number of metrics being combined. (For example, if 4 metrics are being combined, medium sensitivity sites will score 16 and high sensitivity sites will score 81.) Scores are then added across metrics.

The advantage of this system is that if any one metric is scored as high the site is given a higher priority than if all metrics score medium, meaning that it is the most precautionary system. (This is the scoring system IBAT use to rank sensitive sites (IBAT, 2024)).

### 2.8 Comparing identification of priority sites

To examine the influence that the decisions described above have on sites being ranked in the top 20%, we explored;

a. metric choice (based on our basket framework),
b. buffer size,
c. scoring system,
d. sensitivity thresholds.

We used Spearman’s Rank correlation to compare site rankings, Jaccard similarity correlation to compare the identity of sites which fall in the top 20% and rank shift plots to visualise how each site shifts in rank based on metric choice, scoring system and sensitivity threshold.

Metric datasets were overlaid with sites’ AoI with the criteria used to assess the ecological sensitivity of each site based on the maximum metric value found within the AoI (site + buffer). We used each metric to assess the sensitivity of each site using buffer sizes of 50 km (Mining), 10 km (Onshore wind energy and Agriculture) and sensitivity thresholds based on IBAT recommendations for Proximity to PAs, Proximity to KBAs and STAR and our upper quartile system for all other metrics. We then explored site prioritisation as described above. In order to keep the number of comparisons manageable we adopted the following procedures:

1. When testing different metric combinations, we used the Trigger scoring system since it was the most precautionary. To ensure scientific credibility but also simplicity we assigned sensitivity thresholds using IBAT recommendations (for Proximity to PAs, Proximity to KBAs and STAR) and our upper quartile system (all other metrics).
2. When testing different scoring systems, we used the sensitivity thresholds described in the point above and metric combinations A, E and J to ensure every metric was included.
3. When testing different sensitivity thresholds, we used the Trigger scoring system and metric combinations A, E and J.

## 3. Results

### 3.1 How does metric choice influence the identification of ecologically sensitive sites?

As we expected, there is considerable mismatch between sites identified as sensitive by metrics relating to the TNFD’s criteria for ‘areas important for biodiversity’ and ‘areas important for ecosystem integrity’ as well as variation between metrics within these criteria. Whilst every site is identified as sensitive by at least one metric, the percentage of sites identified as sensitive by a single metric, ranges from 2-100% (Figure 3). There is also variation within and between countries in sites identified as sensitive. Biodiverse countries such as Brazil and Indonesia score highly on metrics linked to global extinction risk and species richness. In contrast, the UK, which is recognised as being nature-depleted countries, generally scores low for metrics relating to global extinction risk and ecosystem integrity.

**Figure 3.**
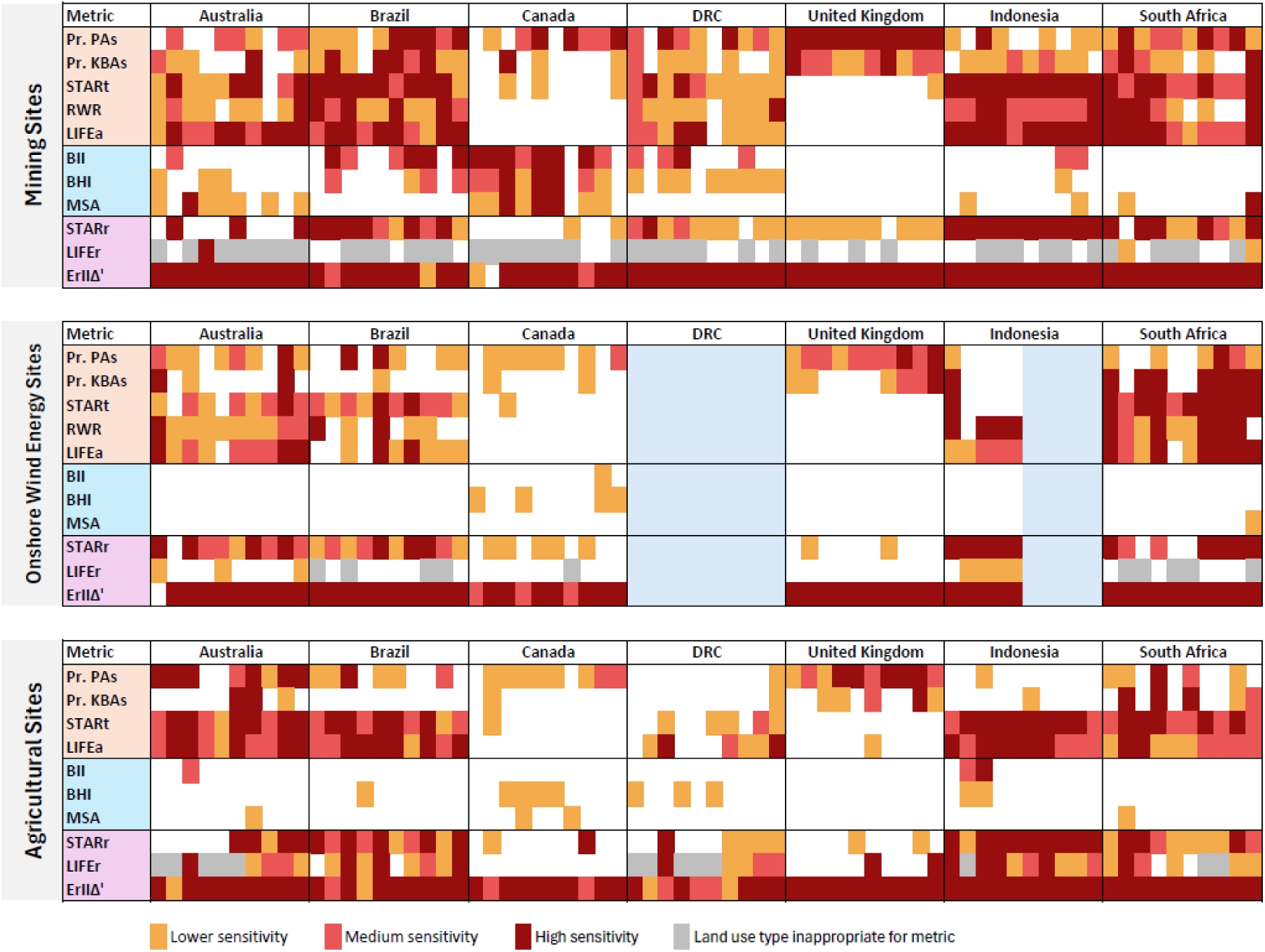
The ecological sensitivity of sites as estimated by different metrics for Mining, Onshore wind energy and Agricultural sites. Site IDs are not shown but run from 1 to 70 (or 55 in the case of onshore wind energy), left to right.

### 3.2 Metric baskets

#### How do the different metric combinations rank sites with respect to ecological sensitivity?

We found that in most cases there was very high correlation (>0.7) across metric combinations with respect to ranking sites for ecological sensitivity (= 0.87, 0.81 and 0.88 for Mining, Onshore wind energy and Agriculture, respectively, Supplementary Information Figure S1). There were particularly high correlations within combinations A-C, D-F and G-J, suggesting the choice between the three ecosystem integrity metrics has little effect on the overall rankings. Correlation scores fall below 0.7 in a few instances for Onshore wind energy, caused by differences between STAR_T_ and RWR.

#### How similar are the top 20% of sites chosen by each combination of metrics with respect to ecological sensitivity?

Our Spearman’s Rank correlation looked at the entire spread of scores from no sensitivity to high sensitivity. However, more relevant for choosing priority sites is an understanding of the similarity of high scoring sites across combinations. Similarity between our Mining sites ranked in the top 20% across combinations ranges from moderate (J >= 0.56) to identical (J = 1). Like the Spearman’s Rank matrix, we see that, within Mining, the greatest area of disagreement comes from combinations A-C and D-F, i.e. between STAR_T_ and RWR (Figure S2). Given the additional threat-related information provided by STAR_T_, we would recommend using STAR_T_ over RWR. Agreement between STAR_T_ and LIFE_A_ is higher in Agricultural sites than in Mining and Onshore wind energy sites which would be expected given that land use change is likely to be the dominant driver of extinction risk in agricultural sites.

#### How frequently are individual sites ranked in the top 20% of ecologically sensitive sites by the different metric combinations?

Following the basket framework, most high-priority sites are identified consistently regardless of metric combination. Within sectors, 50-69% of sites were selected as sensitive by every metric combination (Figure S3). These sites tend to be close to PAs and KBAs meaning that their sensitivity scores are high regardless of how they score on other metrics. Some sites that are identified as sensitive by only a few metric combinations have the same total score across combinations but are assigned different ranks (e.g. Mining Site 19), whilst others score much higher on some metrics than others (e.g. Mining Site 34). More concerning might be cases where sites generally rank highly for sensitivity but are missed by a couple of combinations of metrics. For our sites, however, although there were several instances of sites being identified as sensitive by 6 of the 9 metric combinations, we found no pattern whereby particular metric combinations consistently failed to identify sites as sensitive.

Figure 4 shows the minimum, mean and maximum rank for each site in a rank shift plot. The highest ranked sites (i.e. the most ecologically sensitive) are at the bottom left of the plot and wide bars indicate sites that exhibit high variation in site sensitivity score across the different metric combinations. With respect to prioritising sites at *Locate*, we would be concerned if sites with high mean ranking were exhibiting wide variation in rank across metric combinations as this would be an indication of sites that are ecologically sensitive being missed by a particular metric combination. However, in general, the highest mean ranked sites do not exhibit large differences between their highest and lowest rank. Of Agricultural sites ranking in the top 10 via any metric combination, only 4 have their lowest ranking outside the top 10 (sites 1, 9,10 and 70) and 3 of these do not fall lower than rank 14. However, some sites show high variation in ranking, e.g. site 40 ranks between 11 and 51 due to being identified as highly sensitive by LIFE_A_ and RWR but not sensitive by STAR_T_. The Onshore wind energy sites show a similar pattern with most of the top ranked sites being identified as high priority by all metric combinations. Mining sites exhibit a little more rank shifting amongst high priority sites. In total, 9 sites are given rankings that straddle the top 10 ranking mark and three of these have a wide range of ranking e.g. site 20 ranges from rank 3 to 31 owing to differences in sensitivity scores between STAR_T_, RWR and LIFE_A_. Overall, sites that exhibit greater rank shifting between metric combinations tend to have lower sensitivity scores for proximity to both PAs and KBAs, coupled with variation between their STAR_T_, RWR and LIFE_A_ scores. Sites that are very close to PAs and KBAs are therefore more likely to be identified as ecologically sensitive by all metric combinations.

**Figure 4.**
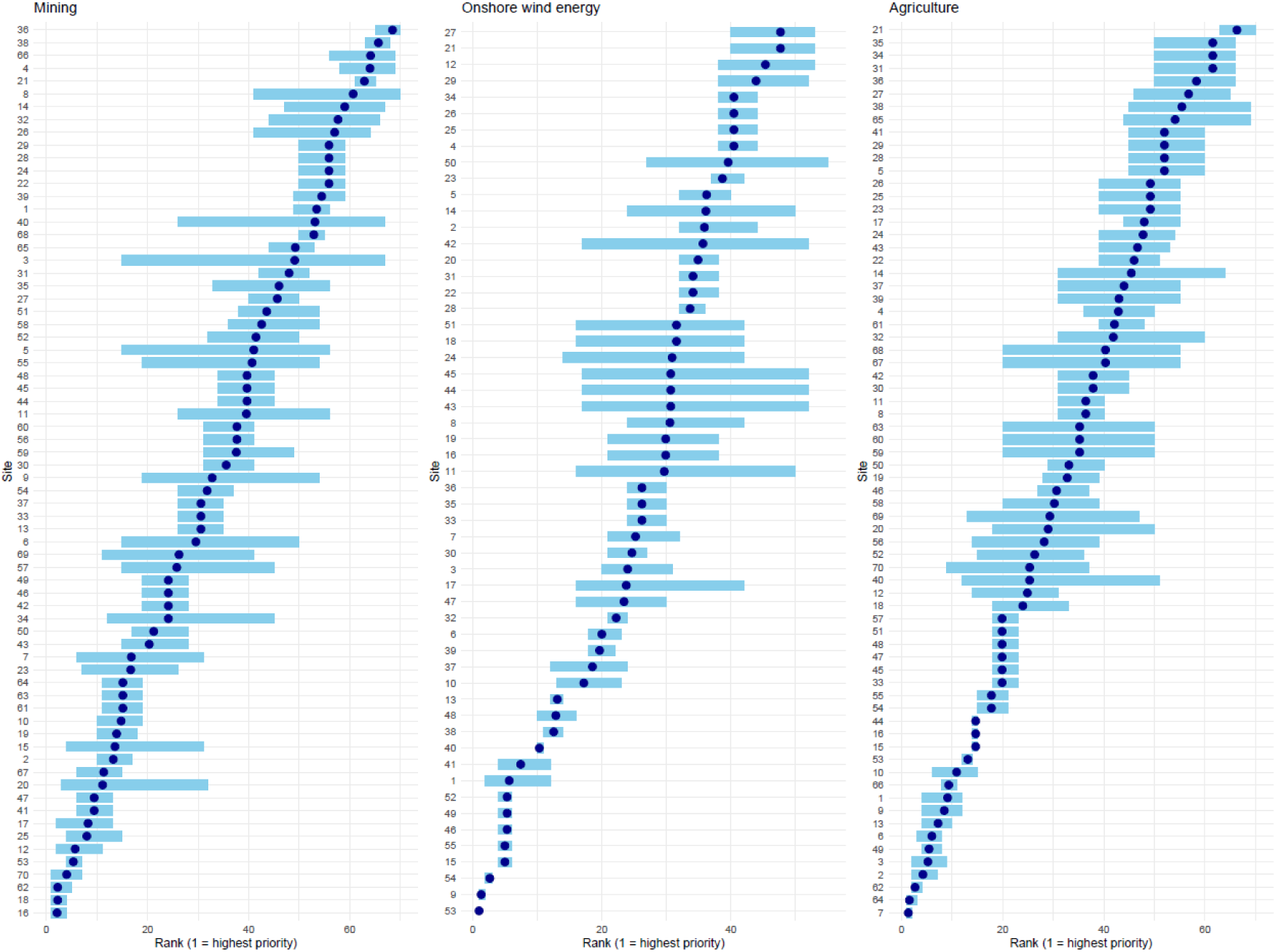
Rank shift plots showing the mean ecological sensitivity ranking (dark blue dot) and maximum and minimum ranking (blue line) for each site based on metric combinations A-J for a) Mining, b) Onshore wind energy and c) Agriculture. (Sensitivity thresholds were based on IBAT recommendations (proximity to PAs, proximity to KBAs, STAR) or the upper quartile definition (all other metrics) and metric scores were combined using the ‘Trigger’ scoring system.)

To summarise, our results suggest that following the basket framework to pick metrics will lead to broadly similar sites being identified as candidates for prioritisation. However, metric choice will have some effect on sites’ rankings, and it is therefore important that companies understand the meaning of the metrics they use.

#### How do the different metric combinations rank sites with respect to restoration value?

The choice of restoration metric led to substantially greater variation in rankings than that caused by species importance metrics. Substituting a restoration-linked metric into the baskets greatly reduced correlation between metric combinations across all sectors. Correlations were as low as 0.42 (Mining), 0.54 (Onshore wind energy) and 0.25 (Agriculture: Figure S4). There were many instances of low similarity in the sites scored in the top 20% with similarity scores as low as 0.27 (Mining), 0.47 (Onshore wind energy) and 0.40 (Agriculture: Figure S5). A lower proportion of sites identified as sensitive by any metric combination (23%, 33% and 31% of sites for Mining, Onshore wind energy and Agriculture respectively) were consistently scored in the top 20% by all K-T combinations (Figure S6). The rank shift plots show that there is reasonable agreement between metric combinations regarding sites consistently ranked in the top 10 for Onshore wind energy and Agriculture (Figure S7). However, there are still wide discrepancies in rankings, particularly in Agriculture, with some sites’ rankings spanning all the way from the top 10 to the bottom 10. Mining sites show considerable variation in rankings with only 3 sites being ranked in the top 10 by all metric combinations.

The choice of restoration metric can thus have a relatively strong influence on the selection of priority sites. These findings suggest that users should consider carefully what they are trying to achieve via restoration before selecting a metric, for example to reduce extinction risk (LIFE_R_ and STAR_R_) or to increase connectivity (ErII Δ’). If users focus on the former, they should consider whether they wish to restrict restoration to arable land (LIFE_R_) or consider all land-use types (STAR_R_).

### 3.3 Area of Influence

#### How does the size of the buffer affect the number of ecologically sensitive sites identified?

As would be expected, for all sectors, the number of sites identified as sensitive increased as buffer size increased (Figure 5). However, the rate of increase differs considerably both between metrics and across sectors. Given the sites we considered represent a very small subset of potential sites globally we can only draw limited conclusions about different sectors, but we can make inferences with respect to sector-specific sites being generally situated in areas of high/low ecosystem integrity and biodiversity importance. Very few of the Onshore wind energy sites were in areas of high ecosystem integrity (Figure 3). Ecosystem metrics associated with ecosystem integrity tended to plateau at low buffer size values for Onshore wind energy sites, i.e. increasing buffer size past the recommended 10 km did not lead to an increase in the number sites being identified as ecologically sensitive. However, this was not the case for metrics identifying the sensitivity of Onshore wind energy sites in relation to biodiversity importance. In the other two sectors, very few metrics plateaued at buffer sizes lower than the IBAT recommendations. We therefore conclude that if buffer sizes below those recommended by IBAT are used, fewer sites will be identified as sensitive.

**Figure 5.**
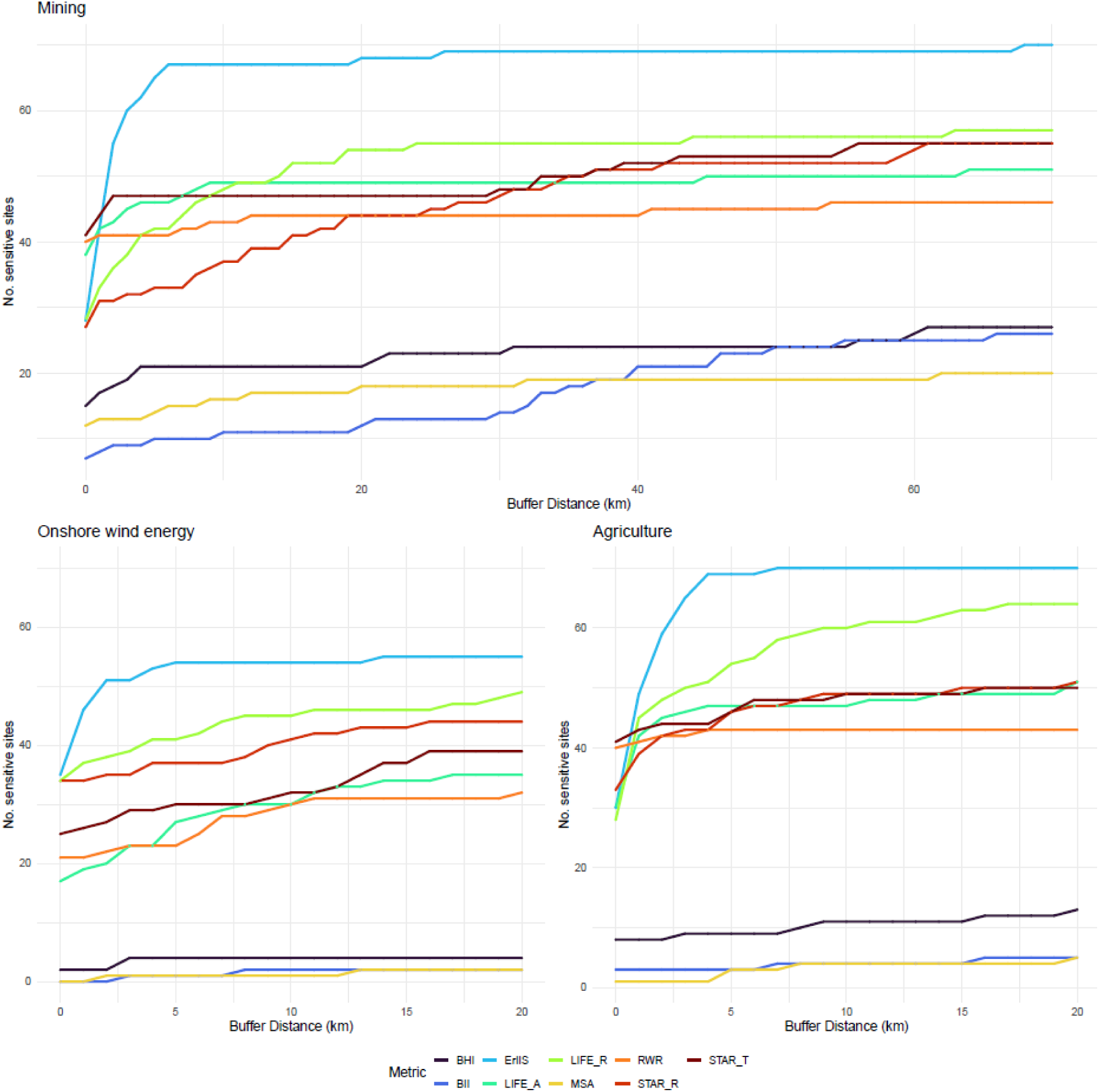
Effect of size of Area of Influence on the number of sites identified as ecologically sensitive for Mining, Onshore wind energy and Agriculture.

#### How does buffer size affect the ranking and selection of ecologically sensitive priority sites?

Site rankings show high correlation between buffer sizes (> 0.7) across all sectors and metric combinations examined (Figure S8). However, similarity values between sites ranked in the top 20% for ecological sensitivity are slightly lower and tend to lie in the moderate range (0.4-0.7) for Mining sites although are still mostly close to or above 0.7 for Onshore wind energy and Agriculture, respectively (Figure S9). In almost all cases, the greatest discrepancy is between the largest and smallest buffer sizes. This difference between Mining and the other two sectors is not unexpected given that the absolute difference in buffer sizes is larger in the Mining sector and that our Mining sites tended to be in areas of higher ecosystem integrity relative to the other two sectors. The majority of Onshore wind energy and Agricultural sites ranked in the top 20% by Combinations A, E and J are selected regardless of buffer size, but this was not always the case in the Mining sector (Figure S10). Nevertheless, we were expecting a greater effect of buffer size on the ranking of sites in the top 20%. The rank shift plot for buffer size shows that sites with a high mean ranking are generally included in the top 20% regardless of buffer size (Figure 6).

**Figure 6.**
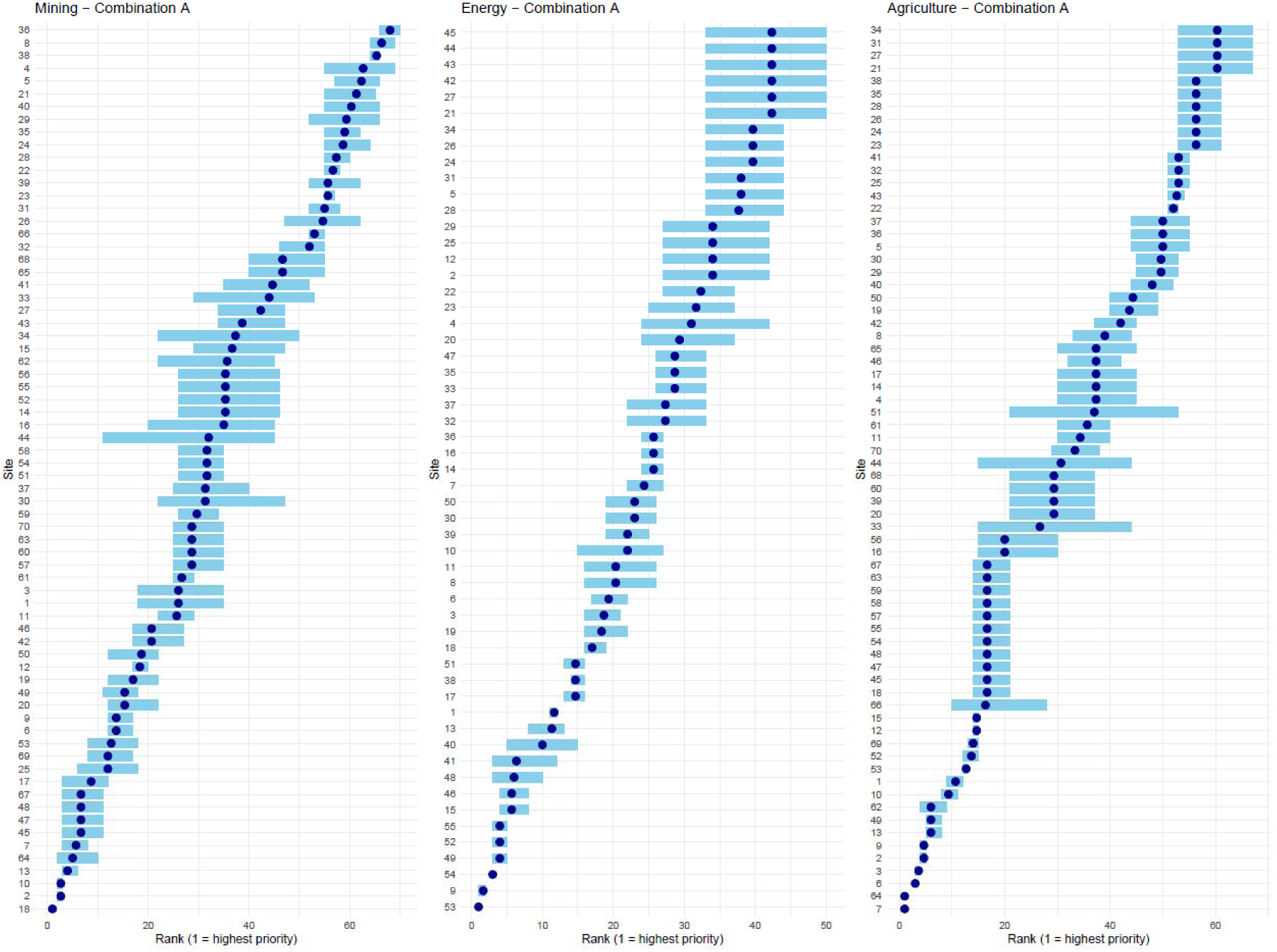
Rank shift plots showing the mean ecological sensitivity ranking (dark blue dot) and maximum and minimum ranking (blue line) for each site for three buffer sizes for metric Combination A for Mining, Onshore wind energy and Agriculture. (Sensitivity thresholds were based on IBAT recommendations (proximity to PAs and KBAs, STAR_t_) or the upper quartile definition (all other metrics) and metrics were combined using the Trigger scoring system.)

### 3.4 Scoring System

#### How do different scoring systems affect the ranking and selection of ecologically sensitive priority sites?

Correlation between different scoring systems for site ranking is high (>0.7) for all three metric combinations considered across all three sectors, particularly so for Onshore wind energy and Agriculture (Figure S12). The lowest correlation is consistently between the Trigger scoring system and the Weighted scoring system. Similarity matrices show that whilst the same sites tend to occur in the top 20% regardless of scoring system for Onshore wind energy and Agriculture (J = 0.69-1.0), scoring system causes large differences in determining which mining sites are ranked in the top 20% (J = 0.27-1) (Figure S13). All four scoring systems rank almost identical sets of sites in the top 20% for Onshore wind energy and Agriculture but this is not the case for Mining for which there is much more variation in site rankings (Figures 7 & S14-15). As noted above, unlike the Mining sites in our analysis, the Onshore wind energy and Agricultural sites are rarely identified as sensitive with respect to ecosystem integrity, which would explain why their ranking is less affected by scoring system.

**Figure 7.**
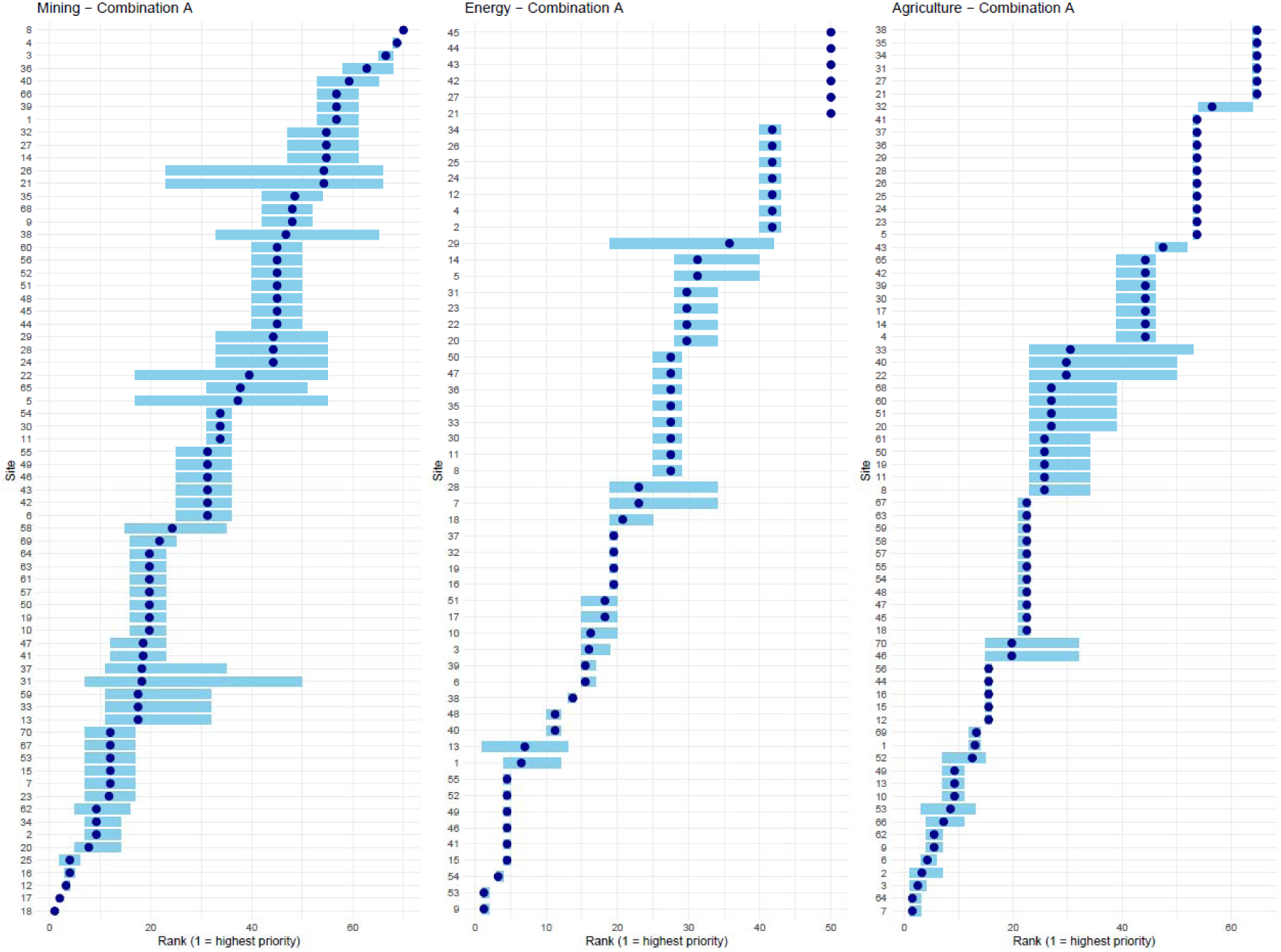
Rank shift plots showing the mean ecological sensitivity ranking (dark blue dot) and maximum and minimum ranking (blue line) for each site for the four scoring systems for metric combination A for Mining, Onshore wind energy and Agriculture. (Sensitivity thresholds were based on IBAT recommendations (proximity to PAs and KBAs, STAR_t_) or the upper quartile definition (all other metrics).)

The Weighted and Multiplicative systems (which both give equal weighting to biodiversity importance and ecosystem integrity metrics) led to moderately similar sites being ranked in the top 20% as did the Additive and Trigger scoring systems (which both give greater weighting to biodiversity importance than ecosystem integrity) but there were large differences between the two sets. These results indicate that more guidance on combining metric scores is required for business. Given that the aim of LEAP’s *Locate* is to identify sites that may be sensitive, we suggest that a precautionary approach of the ‘Trigger’ scoring system may be most useful.

### 3.5 Sensitivity Thresholds

#### How do the different sensitivity thresholds affect the ranking and selection of ecologically sensitive priority sites?

There is moderate to high correlation between the different threshold systems with coefficients ranging from 0.53-0.93 (Figure S16). Metric combination A shows consistently higher correlations across sectors than combinations E and J. There are consistently high correlations (>0.7) between threshold system 1 (50^th^,75^th^,90^th^ percentiles) and threshold system 3 (ecological meaning) for all metric combinations in all sectors. Similarity matrices show that choice of sensitivity threshold can have a large influence over the Mining sites ranked in the top 20% (J = 0.33-0.75) and a low to moderate influence for Onshore wind energy (J = 0.47-1) and Agricultural sites (J = 0.56 – 1) (Figure S17). The different threshold definitions lead to a wider set of sites falling in the top 20% in the Mining sector than within Onshore wind energy and Agriculture (Figure S18). For higher ranked sites, the rank shift plots show a wider variation in ranking with threshold system for Mining sites than for Onshore wind energy and Agriculture (Figures 8 & S19).

**Figure 8.**
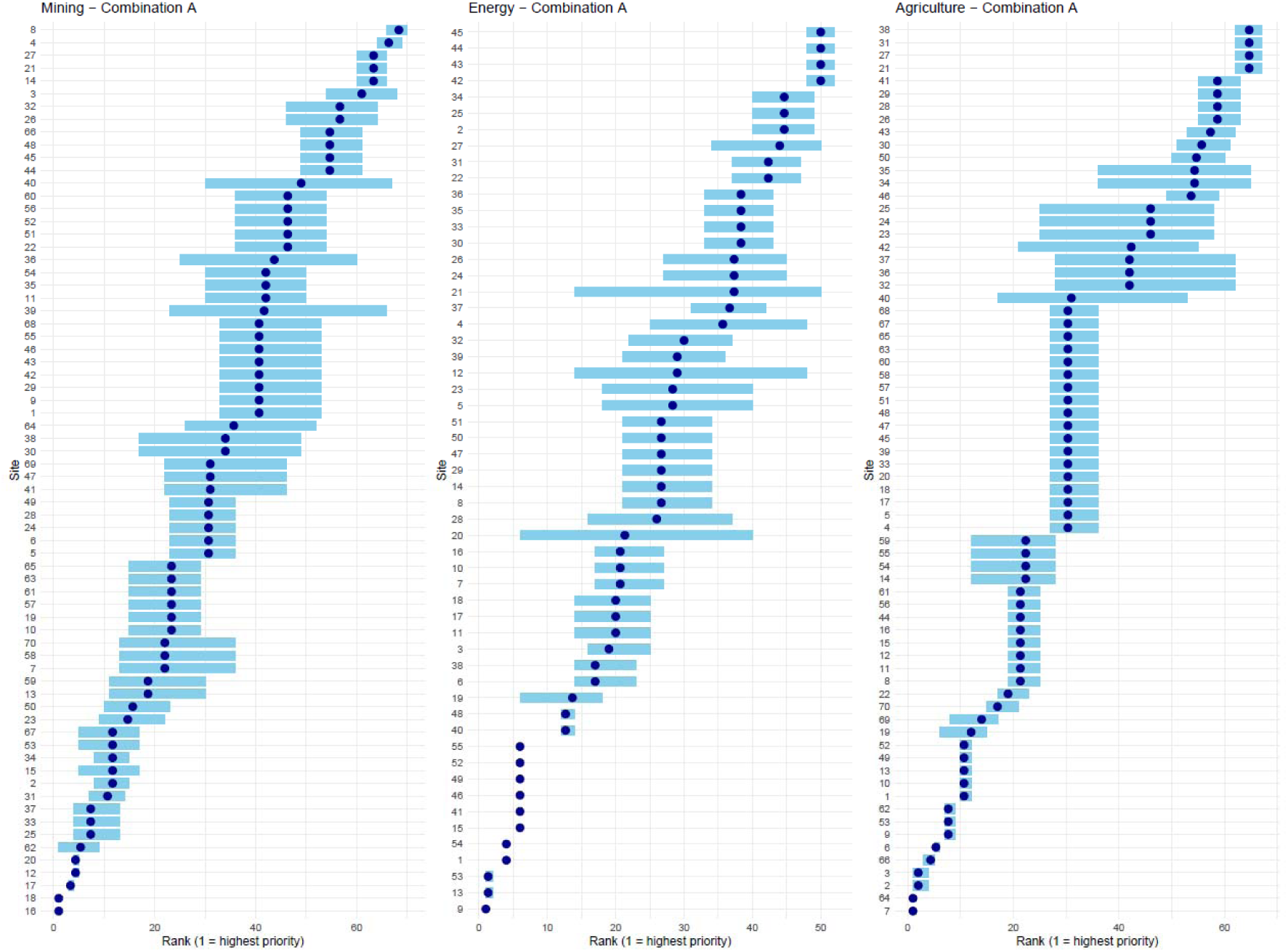
Rank shift plots showing the mean ecological sensitivity ranking (dark blue dot) and maximum and minimum ranking (blue line) for each site for the four sensitivity threshold definitions for metric combinations A, E and J for Mining, Onshore wind energy and Agriculture. (Metrics were combined using the Trigger scoring system.)

Our results show that the choice of sensitivity threshold can have a strong influence over the prioritisation of sites and highlight the importance of organisations being fully transparent in disclosing and justifying their choice of sensitivity thresholds. More guidance on this topic would be helpful for business and would aid disclosure transparency.

## Discussion

We explored the effects of metric choice, buffer size, metric-combination and sensitivity thresholds on the outcomes of assessing sensitive sites in the *Locate* step for nature-related disclosures by business. As would be expected, some metrics scored sites very differently from others but, if metrics relating to biodiversity importance and ecosystem integrity are chosen using our TNFD-based basket framework, site-ranking for the top sites is relatively robust to metric choice (with the exception of screening for sites’ restoration value). However, we found that choices in relation to buffer size, metric scoring system and sensitivity thresholds can have a considerable effect on the set of sites ranked in the top 20% with respect to ecological sensitivity. We suspect that users generally give greater consideration to metric choice than these other aspects, so our results highlight the importance of detailed guidance on these dimensions to help organisations undertake robust nature-related disclosures. Our findings also emphasise the importance of transparency regarding the methods used to locate priority sites when publishing nature-related financial disclosures. When using metrics to locate priority sites for restoration, organisations should choose the metric most appropriate for their purposes, e.g. restoring arable or increasing ecosystem connectivity.

In our analyses, we looked at only two nature-related values (biodiversity importance and ecosystem integrity) but in TNFD’s LEAP process, organisations are also required to assess sites for declines in ecosystem integrity, their importance for ecosystem function and water physical risk. Financial materiality will of course also play a large role in determining priority sites. Given these many considerations, small differences in site rankings caused by methodological choices when screening sites are likely not of great concern providing similar sites are ranked highly.

In our analysis, almost every site was identified as ecologically sensitive by at least one metric. Organisations might be tempted to further filter sites by increasing ecological sensitivity thresholds or decreasing buffer sizes. However, we show that this strategy could lead to nature-related DIROs being overlooked – sensitivity threshold and buffer size can have considerable influence over which sites are ranked in the top 20% so reducing buffer size or increasing sensitivity thresholds risks excluding sites for which companies have high DIROs. It would therefore be preferable to identify all ecologically sensitive sites and then, as a second step, prioritise these sites by, for example, scoring and combining metrics to rank sites in a more structured way.

The TNFD LEAP guidance rightly recognises that there needs to be flexibility for organisations to define buffer sizes and sensitivity thresholds. For example, a site in an area with high pre-existing levels of infrastructure and economic activity will likely have fewer negative impacts on biodiversity than one in a previously roadless area (Ibisch et al., 2016). Likewise, sensitivity thresholds will not have a one-size-fits-all approach. For example, few of our UK sites were identified as sensitive by metrics estimating global extinction risk and ecosystem integrity due to the UK’s low endemicity (i.e. low global extinction risk) and high nature-depletion (i.e. low ecosystem integrity). If an organisation were entirely UK-based, they might therefore want to lower their sensitivity thresholds to identify sites that are of relatively high levels of ecosystem integrity at a national scale. However, there is considerable complexity around understanding the extent of indirect impacts on biodiversity (Narain et al., 2026) and in setting sensitivity thresholds.

At present, none of the existing nature-related disclosure or sustainability standards specify fixed distance buffers to delineate AoI (although the consultation draft for the update to the Sustainability Accounting Standards Board (SASB) Industry-Based Guidance for energy and agriculture proposes a 5 km buffer (International Sustainability Standards Board (ISSB) / IFRS Foundation, 2026)). Instead, the guidance around the different standards states that organisations should choose AoI based on factors such as impact pathway, biome, project characteristics, pollution, hydrology and landscape connectivity (IFC, 2012; SBTN, 2026; TNFD, 2023a). Application of the principles-based standards (e.g. TNFD (TNFD, 2023b), GRI (GRI, 2023), IFC PS6 (IFC, 2012), ESRS (European Commission, 2023)) therefore requires methodological decisions for which there is currently no guidance. To help businesses, IBAT (IBAT, 2026) provides default buffer sizes (with appropriate justifications) but also allows users to apply bespoke buffer sizes. This seems sensible given it is unrealistic to expect organisations to estimate different AoIs for each site when a company’s portfolio may include many sites. Whilst we recognise that a mandatory universal buffer – even if sector-specific – would be scientifically unjustifiable due to the different environments and activities at sites, placing the entire burden on organisations to define AoI feels unworkable given the complexity of the issue and the sparse guidance on estimating indirect impacts in impact assessment practice (Brownlie et al., 2013; Lenzen et al., 2003; Narain et al., 2026). A simple option would be for organisations to begin with the IBAT recommendations for buffer size (IBAT, 2024) and modify these as necessary in conjunction with Narain et al.’s (2026) framework for capturing indirect impacts in site-level screening.

Echoing the various standards’ approach to setting AoI, businesses are also expected to take responsibility for defining ecological sensitivity thresholds for each metric. Unlike climate reporting, there are almost no universally agreed quantitative thresholds for biodiversity metrics other than those proposed by IBAT (IBAT, 2024) and thresholds defined for Critical Habitat as used by IFC PS6 (IFC, 2012). Given there is little scientific consensus on ecological thresholds – indeed it has been suggested that ‘safe-operating spaces’ are likely unquantifiable (Hillebrand et al., 2020), it feels unreasonable to put the onus on business organisations to define their own thresholds. A simple approach for organisations would be to use IBAT’s sensitivity thresholds for proximity to PAs & KBAs and STAR and, in the absence of further guidance for other metrics, or if using global metrics at a national scale, to employ an objective approach such as those we suggest based on quartiles.

Our results showed that the scoring system used to combine metrics can have a considerable influence over which sites are ranked in the top 20%. The Trigger scoring system is the most precautionary – a site scores highly if any individual metric identifies it as highly sensitive. In contrast, a site needs to be sensitive for both biodiversity importance and ecosystem integrity to achieve high rankings using a Multiplicative method. There are arguments in favour of either method, although given the purpose of the *Locate* step, we recommend the Trigger scoring system due to its precautionary nature. It is important that organisations are given guidance to support them to understand the nuance between these methods to make informed decisions over how to combine metrics in line with their specific objectives. Business should be encouraged to disclose these sensitivity findings and provided guidance on how to prioritise further, if a large percentage of the portfolio of sites is deemed high sensitivity.

Objective frameworks and methodologies are essential in assessing nature-related DIROs. However, there is necessarily a degree of subjectivity in the process too, due to, for example, nature’s multi-dimensionality, its lack of fungibility and the spatial variation in its high and often irreplaceable cultural value. Whilst scientific methods can be used to estimate species’ extinction risk or forest cover, a highly threatened species of frog or a sacred stand of ancient woodland may have very different values to different peoples. If both are irreplaceable in some aspect, prioritising one above the other will involve prioritising particular values over others. Similarly, organisations can use objective frameworks to identify sites in ecologically sensitive locations but prioritising among those sites will often involve some level of subjectivity. It is therefore crucial that organisations have a clear understanding of the biodiversity metrics they are using to make informed decisions with respect to interpreting and combining metrics in consideration of multiple stakeholders’ values.

We did not set out to understand how different sectors compare to each other with respect to identifying ecological sensitivity of sites, but to understand better how different methodologies might affect sites chosen at *Locate*. For simplicity, we compared the similarity of highly ranked sites across different sensitivity thresholds using a single scoring system and, likewise, compared scoring system for only one set of sensitivity thresholds. Broader threshold classes may be more stable with respect to ranking and likewise, scoring systems that give less weight to highly sensitive sites may show more similarity across different sensitivity thresholds. Nevertheless, this would not change our main conclusions.

In some circumstances, or for particular sectors, organisations might wish to add in or substitute additional metrics when locating ecologically sensitive sites. The Forest Landscape Integrity Index (FLII), for example, would be particularly relevant for the timber sector or the Global Soil Erosion (GloSEM) dataset for agricultural production. We did not explore these (or other) additional metrics since we would expect their use to lead to differences in the identification and ranking of ecologically sensitive sites. There are several other metrics recommended by the TNFD LEAP approach we also did not use in the analysis either because they remain in development (the World Database of Ecological Corridors (UNEP-WCMC & IUCN, 2026), the IUCN Red List of Ecosystems (IUCN-CEM, 2022), the Ecosystem Integrity Index (Hill et al., 2022)) or because they did not overlap any of our sites (Atlas of Migratory Ungulates (Kauffman et al., 2026)). Our basket framework could be extended in line with the TNFD guidance (TNFD, 2023a) to include baskets to cover metrics associated with the sub-criteria ‘areas containing rare or threatened ecosystems’ and ‘areas important for ecological connectivity’.

In conclusion, the flexible ‘basket’ framework that we developed from the TNFD’s LEAP guidance resulted in similar sites being ranked in the top 20% for ecological sensitivity regardless of metric choice. However, if a business was to use only one or two metrics or to pick metrics in an unstructured way, the identity of sites scored as sensitive would be highly sensitive to metric choice. For our mining company, choices of buffer size, scoring system to combine metrics and ecological sensitivity thresholds all led to moderate to high levels of dissimilarity between top ranking sites and more guidance is needed around these issues, such as frameworks that allow for flexibility but are operable across different nature reporting standards. Clear documentation of assumptions, thresholds and scoring methodologies is essential to support consistency, reproducibility and credibility in TNFD-aligned assessments and disclosures if they are to help protect and restore nature.

## Acknowledgements

This project was supported by the Cambridge Conservation Initiative (CCI) and received research funding from the UBP Nature Finance Initiative and the Royal Society for the Protection of Birds. We thank Emily McKenzie, Rebecca Nohl, Neville Ash, Jacob Bedford, Megan Sim, Mark Leckie, Anne-Sophie Pellier, Melissa Leach, Elizabeth Allen, Victoria Leggett, and Melanie Heath for their interest and support for this work and Alison Eyres, Andy Purvis and Tom Harwood for assistance in reviewing metric summaries.

