## Appendix A for "Exploring the use of state of nature metrics to screen global business operations for ecological sensitivity and to select priority sites for disclosure"

**Appendix A. Summary of Metrics**

The ideal *Locate* metric would measure/be modelled on a good representation (where relevant) of taxonomic groups, terrestrial biomes and geographical regions; be based upon recent data; accurately describe the measurement objective (i.e. areas of biodiversity importance or high ecosystem integrity); be able to be disaggregated to give information on underlying data patterns; inform an organisation’s conservation actions and/or targets; and be applicable at stages of LEAP beyond *Locate*. In reality, no metric will excel across all of these criteria.

In Appendix A we summarise each metric used in our analysis, giving a brief account of key features, underlying data, regularity of updates and associated future plans and limitations.

Appendix B contains spreadsheets detailing the attributes of each metric, the desirable qualities of metrics used to locate ecologically sensitive sites and a summary of how well each metric meets each of those qualities.

### Proximity to Protected Areas (PAs)

*Summary:* The Proximity to Protected Areas (PAs) metric is based on [the [World Database of Protected and Conserved Areas](https://www.protectedplanet.net/en/thematic-areas/wdpa?tab=WDPA)](https://www.protectedplanet.net/en) (UNEP-WCMC and IUCN, 2026) (WDPCA) and can be calculated for sites using [IBAT](https://www.ibat-alliance.org/) (IBAT, 2026).

*Details:* PAs exist under different governance authorities: governance by government, shared governance, private governance and governance by indigenous peoples and local communities. PAs should comply with either the IUCN or Convention for Biodiversity (CBD) definitions:

IUCN definition: “A protected area is a clearly defined geographical space, recognised, dedicated and managed, through legal or other effective means, to achieve the long term conservation of nature with associated ecosystem services and cultural values” (Dudley, 2008).

CBD definition: “A geographically defined area, which is designated or regulated and managed to achieve specific conservation objectives” (Article 2 of the Convention on Biological Diversity (Secretariat of the Convention on Biological Diversity, 2011)). This definition is further expanded upon under Article 8 of the same convention (Secretariat of the Convention on Biological Diversity, 2011).

The WDPCA also contains information on Other Effective Area-Based Conservation Measures (OECMs) which are geographically defined areas outside the PA network that achieve long term and effective in-situ conservation of biodiversity and ecosystem services which may also include cultural, spiritual or socioeconomic values. The OECM network can be viewed as a spatial layer in IBAT (IBAT, 2026) but OECMs are not currently included in IBAT’s Proximity to PA measurement since, to date, they have only been set up in a small number of countries.

*Key features:* Understanding Proximity to PAs is an essential component of environmental due diligence and helps organisations identify both risks and opportunities.

*Underlying data:* A wide range of governmental and non-governmental organisations submit PA data to UNEP-WCMC who manage the [World Database of Protected and Conserved Areas](https://www.protectedplanet.net/en) (UNEP-WCMC and IUCN, 2026).

*Updates*: UNEP-WCMC aims to update all national datasets at a minimum of every 5 years. This is an ongoing process and a new version of the WDPCA is released every month. This means if an organisation was to revisit the *Locate* phase at a later date, they should check whether any new PAs had been designated in the intervening time.

*Limitations:* The vast majority (91%) of PAs are represented by polygons but boundaries are not yet available for the remaining 9% which are represented by points only (UNEP-WCMC, 2019). In these latter instances, although coordinates are requested for the centremost point of the PA, it should not be assumed that the point always represents the centre. Despite formatting and verification standards, the wide variety of data providers and their different capacities and resources mean there are likely to be inaccuracies within the WDPCA database (UNEP-WCMC, 2019). PAs are not necessarily either optimally placed or managed for biodiversity conservation (Pressey et al., 2015).

### Proximity to Key Biodiversity Areas (KBAs)

*Summary:* Proximity to KBAs is based on spatial data from the [World Database of Key Biodiversity Areas](https://www.keybiodiversityareas.org/) (BirdLife International, 2026) (WDKBA) and can be calculated for sites using [IBAT](https://www.ibat-alliance.org/). KBAs are sites of significance for the global persistence of biodiversity.

*Details:* A site qualifies as a global KBA if it meets one or more of 11 specific criteria with quantitative thresholds grouped into 5 higher-level categories relating to: threatened species and ecosystems, geographically restricted species and ecosystems, ecological integrity, biological processes (such as breeding aggregations and recruitment sources), and irreplaceability (IUCN, 2016). The KBA criteria can be applied to species and ecosystems in terrestrial, inland water, marine and subterranean environments, and may be applied across all taxonomic groups. Over 16,500 KBAs have been mapped and documented in virtually all countries and territories worldwide, covering 9.4% of land outside Antarctica and 4.3% of coastal seas (Butchart et al., 2026). KBA assessment is an ongoing process – to date, most KBAs (63%) qualify due to the globally threatened species they support, with 48% being important for biological processes and 39% for geographically restricted species. KBAs have been identified for 17,558 qualifying species in total, of which 36% are plants and 31% are birds (Butchart et al., 2026). The identification of a site as a KBA implies that the site should be managed in ways that ensure the persistence of the biodiversity elements for which it is important. Many KBAs (62%) overlap wholly or partly with existing Protected Areas or Other Effective Area-based Conservation Measures (OECMs) but almost one-third lack any protected coverage (Butchart et al., 2026).

*Key features:* KBAs are widely recognised in various policy processes, with coverage of KBAs by Protected Areas being adopted by the United Nations as official indicators for the Sustainable Development Goals (United Nations, 2024) and being recommended by the Convention on Biological Diversity as part of a headline indicator for the Global Biodiversity Framework (CBD, 2022), as well as adoption or use by other multilateral environmental agreements (Butchart et al., 2026). KBAs are also referenced in various private sector regulations and frameworks, including IFC Performance Standard 6 (IFC, 2012), World Bank’s Environment and Social Standard ESS6 (World Bank, 2017), the Equator Principles (Equator Principles, 2025), various development bank safeguard policies (Butchart et al., 2026), [the European Sustainability Reporting Standards](https://www.unepfi.org/impact/interoperability/european-sustainability-reporting-standards-esrs/) under [the Corporate Sustainability Reporting Directive](https://eur-lex.europa.eu/legal-content/EN/TXT/?uri=CELEX:32022L2464), [the CDP environmental disclosure system](https://www.cdp.net/en), and [the Global Reporting Initiative](https://www.globalreporting.org/) (Butchart et al., 2026).

*Underlying data:* The World Database of Key Biodiversity Areas (WDKBA) is managed by BirdLife International on behalf of the KBA Partnership which comprises 13 of the world’s leading nature conservation organisations including IUCN, BirdLife International and RSPB.

*Updates:* The KBA guidelines state that sites should be reassessed at a minimum of every 8-12 years (KBA Standards and Appeals Committee of IUCN SSC/WCPA, 2022).

*Future Plans:* Comprehensive KBA assessments are underway in many countries around the world, which is rapidly growing the number of sites identified for lesser-known vertebrate groups, invertebrates, plants and fungi, including in freshwater, marine and subterranean environments.

*Limitations:* Although KBAs have been identified for mammals, birds, amphibians, reptiles, fish, plants and invertebrates (mostly Insecta, Gastropoda, Anthozoa and Malacostraca), there are fewer sites identified for invertebrates – to date, 36% of qualifying species are plants and 31% are birds (KBA Programme, 2026). Fewer sites have been identified so far under the criteria relating to ecosystems, ecological integrity and irreplaceability than those relating to species, also relating to knowledge gaps (e.g. risk of ecosystem collapse has not yet been assessed at a relevant level of resolution globally) (Butchart et al., 2026). There are still many sites which likely fulfil the criteria of a KBA but that have yet to be formally identified (KBA Programme, 2026).

### Species threat abatement and restoration (STAR)

*Summary:* STAR evaluates the potential contribution to reducing global extinction risk of species from actions to reduce threats and restore habitat at particular locations. STAR has two complementary elements, STAR for Threat Abatement (STAR_T_) and STAR for Restoration (STAR_R_) (Mair et al., 2021). STAR scores are available through IBAT (IBAT, 2026). STAR_T_ scores can be broken down into scores for specific threats facing individual species. This enables identification of targeted actions that could be taken to address those threats.

*Details:* Areas with high STAR_T_ contain higher numbers of more highly threatened species and larger proportions of individual species’ ranges. These are areas where interventions could make a large contribution to reducing global species extinction risk. Indonesia, Colombia, Mexico, Madagascar and Brazil together contribute to over 30% of the global STAR_T_ score (Mair et al., 2021). STAR_T_ scores can be broken down to show the contributions of different threat types, e.g. invasive species, pollution, logging and wood harvesting.

Areas with high STAR_R_ represent locations which previously supported higher numbers of more highly threatened species and larger proportions of individual species’ ranges. These are locations where historical impacts have occurred, and restoration would provide an opportunity to reduce species extinction risk. A calculation is built into STAR_R_ scores to reflect the slower and lower success rate in reducing extinction risk by restoring habitat compared with conserving existing habitat.

*Key features:* STAR supports the establishment of science-based targets across spatial scales, because it enables the potential impacts of different actions in different locations for different species to be quantified in a comparable way and aggregated. It can therefore be used at every stage of the LEAP process – see [The IUCN RHINO Technical Source Document](https://cdn.prod.website-files.com/68b5bcab2e9b175c9bf0fa8e/68e7bd96fea70908ba9b51f6_IUCN%20RHINO%20Technical%20Source%20Doc%20V2.pdf) (IUCN, 2025a) for further guidance.

*Underlying data:* Species extinction risk categories, threat classification data and range maps are from the IUCN Red List (IUCN, 2025b). Area of Habitat maps were calculated using IUCN range maps (IUCN, 2025b), the European Space Agency Climate Change Initiative (ESA CCI) land use and cover maps (ESA, 2017) and Copernicus Global Land Service 100m Land Cover maps (Buchhorn et al., 2020).

*Updates*: STAR will be updated with each issue of the IUCN Red List (currently twice yearly) to reflect the latest information on taxonomy, species’ extinction risk, habitat preferences, elevational limits, distribution and land cover. However, due to the enormous amount of work involved, it is important to note that not all species are reassessed with each Red List update. Note that sensitivity thresholds are currently based on global median values of STAR and could change – refer to IBAT (IBAT, 2026) for updated thresholds.

*Future plans:* STAR is currently based on data for mammals, birds, amphibians and reptiles and there are plans to add further taxa such as plants. It is currently only available for terrestrial species, but a combined terrestrial, freshwater and marine layer is in development. IBAT will shortly be adding a feature to calculate an updated Calibrated STAR score (Mair et al., 2026) following ground-truthing of species and threat data at priority sites.

*Limitations*: STAR treats species classified as Least Concern as having zero global extinction risk. A species may not be present everywhere within its mapped Area of Habitat and threats may not impact a species equally across its distribution. Hence, when using STAR to set targets and monitor their achievement, it is important to calibrate STAR before implementing actions and monitoring their impact to generate realised STAR values (IUCN, 2025a).

### Rarity-Weighted Richness

*Summary:* Rarity-weighted richness shows the relative importance of an area in terms of its aggregate contribution to the global distribution of species. The measure was developed as a way of weighting species richness by species rarity.

*Details:* Each species in each grid cell is assigned a score calculated by dividing the area of a grid cell by the total area of that species’ geographic range. Scores are then summed across all species present in a grid cell. Higher values show that a cell holds more species and/or a higher proportion of their global ranges. Rarity-weighted richness tends to be higher in the tropics than in temperate regions. This is because the tropics support a) a greater number of species and b) high levels of endemism, i.e. species are more likely to be restricted to a single geographic location, e.g. island or mountain range. In contrast, there is lower species richness in temperate regions and species generally have much wider distributions.

Rarity-weighted richness can be disaggregated by species groups and is available from the IUCN in various different formats with raster layers calculated for specific taxonomic groups (e.g. Birds, Mammals, Birds & Mammals, All species, Threatened species etc) (IUCN, 2025b).

*Key features:* Conceptually, rarity-weighted richness and STAR_T_ are closely related: the global STAR_T_ layer is effectively rarity-weighted richness weighted by extinction risk. However, the data associated with STAR_T_ are informative about the relative contribution of different threats in a particular location to extinction risk, and we would therefore recommend using STAR over rarity-weighted richness since it is more informative.

*Underlying data:* The IUCN has calculated rarity-weighted richness using two different types of input data: range map polygons and Area of Habitat (AoH) rasters. Range maps show the distributional limits for each species, including unsuitable habitat and unoccupied areas (their purpose is to enable the estimation of the spread of extinction risk, not the occupancy of range) The range polygon rarity-weighted richness datasets are based on the Red List version 2-25-2. AoH rasters represent the distribution of suitable habitat types within the elevation limits of the species and are currently available for birds and mammals (IUCN, 2025b; Lumbierres et al., 2022).

*Future Plans*: Area of Habitat maps are being developed for all species for which Red List assessments, range maps and habitat coding are available.

*Limitations:* The current rarity-weighted richness layer range is restricted to birds and mammals, but will shortly be available for all species on the Red List with mapped ranges and distribution information (>100,000 species). Although the species distribution maps underlying rarity-weighted richness are based on recent versions of the Red List, the underlying data, including species occurrences, may derive from a range of dates. Use of this layer to identify species potentially present in an Area of Influence will therefore require ground-truthing.

### Land-cover Change Impacts on Future Extinctions (LIFE)

*Summary:* LIFE estimates changes in the expected number of extinctions (both increases and decreases) caused by converting remaining natural vegetation to arable land (LIFE_convert_arable (LIFE_A_)) and restoring both arable and pasture to natural habitat (LIFE_restore (LIFE_R_)) (Eyres et al., 2025).

*Details:* LIFE uses changes in species’ Area of Habitat (AoH) as a proxy for extinction risk. Estimates of extinction risk are derived from estimates of a species’ current AoH and of its potential AoH in the absence of people. LIFE estimates the change in extinction probability resulting from present day land cover change manifested over the next 100 years. It is assumed that extinction risk responds non-linearly to habitat loss i.e. that a 100km^2^ loss of habitat for a species currently occupying 1000km^2^ will have a greater impact than for a species occupying 1 million km^2^. For each species, LIFE calculates the change in its probability of extinction (relative to that in the absence of human influence) that would occur following the change in the species’ AoH as a result of one unit of land conversion. This procedure is carried out for all species. The final LIFE score for each 1 arc minute grid cell is the estimated global impact of changing land cover (either conversion or restoration) taken as the mean across all species per km^2^ of land use under transition. Extending this across all grid cells produces a global map for each land cover transition scenario. To date, LIFE layers have been produced for two key land cover changes: in calculating LIFE_A_, a scenario is developed in which all terrestrial habitats mapped as non-urban are converted to arable land; for LIFE_R_, the scenario restores all areas classified as arable or pasture to their potential natural vegetation. Species which currently have a population size greater than that of their original (i.e. which have benefited from human activity) only contribute to LIFE scores if land cover changes results in their AoH dropping below the original population size. Land classified as urban is excluded as it is unlikely to change (Eyres et al., 2025).

LIFE covers 30,875 species of terrestrial vertebrates at a 1 arc-min resolution (3.4sq km at equator) but could be used at scales of 0.5 -1000 sq-km. High LIFE scores correspond to cells with higher species richness, cells with a greater than average number of endemic species and/or cells whose species have already lost more of their original AoH (Eyres et al., 2025).

*Key features:* LIFE covers non-threatened as well as threatened species. LIFE takes into account the cumulative negative impact of habitat loss over the long term – most other metrics assume the loss of a fixed area of habitat will have the same effect on a species regardless of its original AoH.

*Underlying data*: AoH maps were calculated from the IUCN Red List’s range maps, habitat preferences and elevational range (IUCN, 2025b), the Global Map of Terrestrial Habitat Types (Jung et al., 2020) (dating from 2016) and the Potential Natural Vegetation map (Hengl et al., 2020).

*Future plans:* Whilst LIFE originally focused on conversion to or restoration of agricultural land, scores for other relevant land use types have been produced, e.g. conversion to pasture or urban. LIFE was originally designed to focus on land cover change (a major driver of biodiversity loss). Current work is focussing on incorporating other threats, e.g. hunting, and on increasing the taxonomic scope of the metric.

*Limitations:* LIFE scores are derived for vertebrates only and should be treated cautiously in regions with particularly high richness and endemism among non-vertebrate groups (e.g. Mediterranean biomes and the Cerrado) (Eyres et al., 2025). LIFE is particularly relevant in scenario modelling, estimating the impact on extinctions of diverse actions that affect change in land cover, for example, dietary choices and protected area development. Within the context of the TNFD, LIFE would likely be most relevant in assessing decisions regarding conversion/restoration of agricultural land.

### The Biodiversity Intactness Index (BII)

*Summary:* The Biodiversity Intactness Index (BII) gives the percentage of the original number of species and their abundance estimated to remain in any given area, despite human impacts (Scholes & Biggs, 2005). BII can be calculated from direct observations at sites or it can be modelled. The two ways that the ‘BII’ term is used – (i) a metric based on direct observations and (ii) a modelled map layer - can cause considerable confusion, even amongst the scientific community. At Locate, BII is used in the context of a modelled global map layer since it would be far too resource intensive to calculate BII from direct observations. (Even at the Evaluate stage calculating BII from direct observations would most likely be impractical.) Further confusion is brought about by different research groups and consultancies using slightly different methods and data to estimate BII.

*Details:* One of the most well-used models of global BII is developed by a research group based at the Natural History Museum, London (De Palma et al., 2024). Their model estimates the average state of local biodiversity via the PREDICTS database (Hudson et al., 2016) which is used to model the response of biodiversity to land-use, land-use intensity, human population density and related pressures. However, consultancies may produce their own models of BII and BII’s analytical approach can be used on geographical, ecological or taxonomic subsets of the underlying PREDICTS database in order to more accurately model specific scenarios, for example, evaluating BII for particular biomes or regions, or evaluating the impacts of specific crop types or other particular land usages.

In our Walkthrough, we used the Natural History Museum’s version 2.1.1 of BII and this is the model we now describe. The global BII map version 2.1.1 is underpinned by the PREDICTS database (Hudson et al., 2016) which contains data on over 60,000 species with a high representation of plant and insect species as well as covering vertebrates and fungi and representing threatened and non-threatened species. BII models the response of biodiversity to six core land-use classes (Primary, Secondary, Cropland, Plantation Forest, Pasture and Urban) at three use-intensity classes (low, medium or high). Agricultural land use is resolved more finely where possible into Annual crops, Perennial crops, Timber plantations, Rangeland and Managed Pasture. The PREDICTS database allows comparisons to be made of the number and diversity of species found within these different land-use classes to those at near-undisturbed sites. These changes in biodiversity can then be averaged across the different land-uses to produce predictions of the effect of a particular type of land-use change on biodiversity. A global map of BII is calculated by combining these predictions with models of land-use derived from satellite imagery (De Palma et al., 2024).

BII has been used in the planetary boundary framework with a precautionary boundary set at 90% (Steffen et al., 2015) (although there is considerable uncertainty around this with the true safe limit estimated to be somewhere between 90-30%.) Crossing the planetary boundary increases the risk that the ecosystem’s functionality is impaired and it can no longer provide key services such as carbon storage or clean air and water. Ecosystem collapse is likely to occur if BII is reduced to below 30% (Newbold et al., 2015).

The methodology behind BII produced by other consultancies may differ from that described above and will not be based on the full PREDICTS database.

*Key features:* BII models based on the full PREDICTS database have excellent representation of species types covering vertebrates, invertebrates, plants and fungi. Models also allow for different land use intensities. BII can be used to model future scenarios at further stages of LEAP. Whilst BII scores cannot be disaggregated into species types, BII scores can be developed for particular regions and/or crop types, increasing accuracy. The Natural History Museum’s BII is the only version of BII which is modelled on the full PREDICTS database.

*Underlying data for the Natural History Museum’s version 2.1.1:* The PREDICTS database (Hudson et al., 2016), CSIRO land-use map (based on ESA-CCI Land Cover data (ESA, 2017), LUH2 Land use layers (Hurtt et al., 2020) and MODIS v6.1 VCF (DiMiceli et al., 2022)), CEISIN human population density data (CEISIN, 2018), GISD30 impermeable surface data (Zhang et al., 2022).

*Updates:* The land-use BII is continually being refined via increasing the data that it draws from and adjusting models for specific scenarios. The Natural History Museum anticipate providing updates at least annually. A 1km resolution dataset is currently released annually to subscribers. A 10-km resolution dataset is available for non-commercial use and is updated approximately every ten years. Contact the Natural History Museum for the latest developments.

*Future plans:* The Natural History Museum plan to improve the representation of land use intensity within the BII pipeline and, subject to funding, to incorporate a wider range of anthropogenic pressures and mitigations to provide a more comprehensive assessment of biodiversity change. They are also intending to produce 1km resolution projections of BII from 2025 to 2050 under a suite of future scenarios (the Shared Socioeconomic Pathway scenarios) during 2026.

*Limitations:* BII measures one component of ecosystem integrity only – composition – and does not cover structural or functional integrity. The global map of BII is based on a model so it is predictive, not observational - it should be ground-truthed at priority sites. Global BII should be used with some caution in areas with plantation forest, particularly intensive plantation, since the underlying land-use layer (in common with most global land-use layers) does not separate plantation from other forests. The global BII layer used in this case-study groups plantation forests with secondary vegetation (De Palma et al., 2024). The pressure-response relationships used in the underlying PREDICTS model assume that biodiversity’s response within individual land use classes is consistent across the globe; in reality, biodiversity’s sensitivity to land use change differs, for example, across biomes and with proximity to highly intact wilderness habitat.

### Mean Species Abundance (MSA)

*Summary:* MSA is an indicator of local biodiversity intactness. It compares the abundance of individual species under the influence of a given human pressure to their abundance in an undisturbed situation (Schipper et al., 2020). MSA ranges from 0 to 1, where 1 indicates a fully intact species assemblage and 0 means that all original species are locally extinct. MSA could in theory be calculated from direct observations at sites but at the Locate stage this would be far too resource intensive. Instead, a global map of MSA estimating the average state of local biodiversity is estimated using the GLOBIO4 model (Schipper et al., 2020). The two ways that the ‘MSA’ term is used – (i) a metric based on direct observations and (ii) a modelled map layer - can cause considerable confusion, even amongst the scientific community.

*Details:* The global MSA map layer is underpinned by GLOBIO4 which models species’ responses to six human pressures: land use, road disturbance, fragmentation, hunting, atmospheric nitrogen deposition and climate change (Schipper et al., 2020). Species’ response to land use is modelled for six land use classes: primary, secondary, urban, cropland, pasture and forestry with cropland and pasture differentiated into two use-intensity classes. The pressures are modelled independently of each other and do not allow for interactions between the pressures (e.g. climate change exacerbating the effects of land use change) although for some land use classes other pressures are ignored as the land use type is assumed to dominate, e.g. road disturbance does not add further biodiversity loss to that estimated for urban land use. Responses to climate change are modelled separately for tropical and non-tropical regions. Models are based on mammals, birds and plants and cover both threatened and non-threatened species. MSA ignores any increases in species above that of the undisturbed situation meaning that the metric is not influenced by species which may benefit from habitat disturbance or climate change, for example.

The global MSA map layer is at a spatial resolution of 300m x 300m. MSA can be combined with the extent (km^2^) of the ecosystem measured to calculate a condition-weighted area (MSA.km^2^).

As explained above, the MSA metric can be calculated based on detailed species abundance field observations and so it could, in theory, be used at the *Evaluate* and *Prepare* stages to assess impact and set targets although this would almost certainly be too resource-intensive in practice. MSA as estimated by GLOBIO4, should not be used to assess fine-scale impact or set targets since it is modelled not directly observed.

*Key features*: MSA (GLOBIO4) excludes non-native species and species which increase in abundance due to human disturbance. MSA (GLOBIO4) models the impacts of multiple anthropogenic drivers of biodiversity change - land use, road disturbance, fragmentation, hunting, atmospheric nitrogen deposition and climate change – and combines them into a single score. GLOBIO4 projects MSA from 2015 to 2050 using three different scenarios meaning it can be used in assessing future risk.

*Underlying data:* GLOBIO4 is underpinned by many different datasets. For simplicity, we list the core datasets only: a sample of the PREDICTS database (Hudson et al., 2016) – species' response to land use and to habitat fragmentation; Nunez et al., 2019 – species' response to climate change; Midolo et al., 2019 – species' response to atmospheric nitrogen deposition; Benítez-López et al., 2010 - species’ response to road disturbance; Benítez-López et al., 2017 & Benítez-López et al., 2019 - species’ response to hunting; ESA-CCI Land Cover data (ESA, 2017), LUH2 Land use layers (Hurtt et al., 2020) and GRIP database (Meijer et al., 2018) – land use pressures.

*Limitations*: MSA measures one component of ecosystem integrity only – composition – and does not cover structural or functional integrity. It is important to understand that the global map of MSA is modelled and is therefore predictive, not observational – it should be ground-truthed at priority sites. The GLOBIO pressure-impact relationships which underpin the global MSA map assume biodiversity’s response within individual land use classes is consistent across the globe; in reality, biodiversity’s sensitivity to land use change differs, for example, across biomes and with proximity to highly intact wilderness habitat. The pressure-impact relationships are based on relatively few values (around 70 sites for land use for example, in comparison to >5000 sites used in BII v2.1.1) and there are significant geographic gaps in their coverage – for animals, for example, there are no land use MSA estimates from Europe and only two in North America meaning estimates of global responses will primarily be based on data from the Tropics. The land-cover map used in the model does not distinguish between primary and secondary vegetation.

### Biodiversity Habitat Index (BHI)

*Summary:* The Biodiversity Habitat Index (BHI) estimates the proportion of species diversity retained within any given area in relation to the degree of habitat loss, degradation and fragmentation of that area (Hoskins et al., 2020). BHI ranges from 0 (total loss of habitat and thus species) to 1 (no habitat degradation and thus no loss of species) and is based on the integration of habitat condition derived from remotely-sensed land cover data and a suite of spatially variable models of the effect of environment on species. Results for the indicator are expressed as the proportion of species in the reporting unit (i.e. grid cell) expected to persist over the long term.

*Details:* BHI can be calculated for any given spatial reporting unit, e.g. a grid cell, a country or a broad ecosystem type but for business use there would be no need for this aggregation. The ‘off the shelf’ version of BHI available for direct download ([CSIRO BHI v4](https://data.csiro.au/collection/csiro:72585v4)) is a raster with 30 arc-second resolution.

BHI is generated from two main inputs – (i) data indicating the present condition or integrity of the natural ecosystem associated with each cell and (ii) pre-derived modelling of spatial variation in the species composition of ecological communities as a function of different environmental conditions (Hoskins et al., 2020). These two inputs are combined to assess the ‘effective proportion of habitat’ which is ecologically similar to all other cells, adjusting for the effects of the condition and functional connectivity of habitat and of spatial variation in the species composition of ecological communities. This ‘effective proportion of habitat’ can then be translated into a prediction of the proportion of species expected to persist over the long term.

BHI is calculated using the Commonwealth Scientific and Industrial Research Organisation’s (CSIRO) BILBI modelling infrastructure (Hoskins et al., 2020). BILBI is based on a global 30 arc-second grid, with environmental data comprising terrain adjusted climate data and soil data from www.SoilGrids.org (v1). This is combined with species data from the Global Biodiversity Information Facility (GBIF) (>400,000 plant, vertebrate and invertebrate species from >300 million location records) to generate spatial biodiversity models (using a technique called Generalised Dissimilarity Modelling (GDM)) for each major biogeographical realm. Downscaled land use mapping estimated from remotely-sensed land-cover change and environmental modelling, is combined with coefficients from the [PREDICTS database](https://www.nhm.ac.uk/our-science/research/projects/predicts.html) (Hudson et al., 2016) which estimate change in species richness and abundance following land use change to produce a global map of the condition of habitat in units of effective cell area of undisturbed native habitat. (As an area of native vegetation is reduced, the number of species it can support reduces, following a predictable curve known as the species-area relationship. The effective cell area of undisturbed native habitat is derived from this relationship and can be thought of as the hypothetical area of native vegetation (expressed as a proportion of the cell’s total area) that would be needed to support the current number of species estimated to occur in the cell.) The raw condition layers are converted into an area-scaled measurement of connected condition using a least cost path model (Drielsma et al., 2007). The connected habitat condition layers are then integrated with the spatial biodiversity models to generate the BHI.

*Key features:* BHI uses coefficients derived from the PREDICTS database (PREDICTS has good representation of terrestrial invertebrates, plants and fungi as well as vertebrates – see BII details) to estimate the impact of land use change on species (represented in BHI by terrestrial plants, vertebrates and invertebrates), but also integrates habitat connectivity – areas with a given level of disturbance will vary in their species composition dependent on their proximity to large regions of intact habitat. BHI estimates are available at annual intervals from 2000 through to 2024 meaning that the index can be used to estimate recent rates of change in ecosystem integrity. BHI’s analytical approach can be used on geographic subsets of data, e.g. national ecosystem condition, to create more accurate local models but computing BHI is extremely computationally expensive and this would likely not be practical for disclosure purposes. BHI could also be used to predict the impacts of proposed actions but, again, this would require considerable expertise and computational time.

*Underlying data*: CSIRO BILBI biodiversity modelling infrastructure based on W[orldClim](https://www.WorldClim) (Fick & Hijmans, 2017), SoilGrids (Poggio et al., 2021), GBIF and RESOLVE ecoregions (Dinerstein et al., 2017) – GDM models; ESA –CSI Land cover (ESA, 2017) and MODIS VCF (DiMiceli et al., 2022) – land cover data; Global Tree Cover 2000 (Hansen et al., 2013); PREDICTS database (Hudson et al., 2016) – coefficients describing species’ response to land use change. Note that the raw condition layer is very similar to that used in BII and the connected condition layer similar to that used in FLII.

*Updates:* BHI is available at yearly intervals from 2000 to 2024.

*Limitations:* The habitat condition surface is derived from a combination of remotely sensed data (which may have errors) and land use specific coefficients from the PREDICTS database. Due to the availability of studies within the PREDICTS database, these coefficients assume a globally consistent effect of anthropogenic land use on the capacity of a location to support natural systems; in reality, biodiversity’s sensitivity to land use change differs across biomes.

The compositional models which are used to build the BHI from the condition surface do not represent the structural or functional components of ecosystem integrity. The global map of BHI is based on a model so the response to cellwise habitat condition is predictive, not observational. It also represents the long term implications of land use change rather than changes that might be seen immediately. The GDM models are currently implemented at the Realm level (e.g. Afrotropic, Neotropic, Indo-Malay), to reflect the differing response of each realm’s biodiversity to the effects of environmental change; in reality, responses will differ across biomes within realms.

### Ecoregion Intactness Index (ErII)

*Summary:* The Ecoregion Intactness Index (ErII) estimates the combined impact of habitat loss, fragmentation and degradation arising from anthropogenic disturbance to give an intactness score for each of the world’s terrestrial ecoregions ranging from 0 to 100 (Beyer et al., 2020). ErII uses the Human Footprint Index (Venter et al., 2016) as a proxy for habitat quality, transforming footprint scores to habitat quality scores.

*Details:* ErII contains three datasets – a polygon layer giving a relative intactness value for each ecoregion and two raster layers, Q’ and delta Q’, which are relevant to the prioritisation of management actions (Beyer et al., 2020). Q’ describes the contribution of each cell to intactness with high values indicating areas that are most in need of protection in order to maintain intactness and ranges from 0 to 1. Delta Q’ measures the expected rate of change in the contribution of a cell to intactness as the quality of habitat in the cell increases or decreases. High values of Delta Q’ indicate areas where habitat rehabilitation is expected to have the largest impact on intactness. Delta Q’ is generally low in regions dominated by low intactness, modest in areas dominated by high intactness, and highest in areas with intermediate levels of intactness.

*Caution:* It is easy to confuse the Ecoregion Intactness Index with the Ecological Integrity Index(Hill et al., 2022)! We have followed the TNFD in abbreviating the Ecoregion Intactness Index to ‘ErII’ to differentiate it from the Ecosystem Integrity Index or ‘EII’, an additional index mentioned in the LEAP guidance. (We have not included EII in our walkthrough examples since it is still under development.)

*Key features:* ErII estimates the combined impact on ecosystem integrity of multiple different anthropogenic stressors: built environments, population density, electric infrastructure, croplands, pasture, roads, railways and navigable waterways. The metric is standardised meaning it is comparable across different ecoregions, whatever their shape or area.

*Underlying data:* The Human Footprint Index (Venter et al., 2016), RESOLVE Ecoregions 2017 (Dinerstein et al., 2017).

*Limitations:* ErII represents the intactness of ecosystems but does not explicitly measure ecosystem integrity by either structure, composition or function. The global map of ErII is based on a model so it is predictive, not observational - it should be ground-truthed at priority sites. Ecoregions smaller than 100km^2^ are excluded because they cannot be characterised at the mapping resolution of the Human Footprint Index. The model assumes that the benefit provided by action within a cell is independent of actions in other cells. The Human Footprint Index may not reflect all anthropogenic pressures such as those that are intermittent (e.g. selective harvesting of forests) or have nonlocal effects (e.g. pollution or pathogens) (Beyer et al., 2020). ErII is not based on recent data – it uses a version of the Human Footprint Index relating to 2009. The Human Footprint Index is not always a good proxy for the value of biodiversity habitat (Mokany et al., 2020).
