## Supplementary Figures for "Exploring the use of state of nature metrics to screen global business operations for ecological sensitivity and to select priority sites for disclosure"

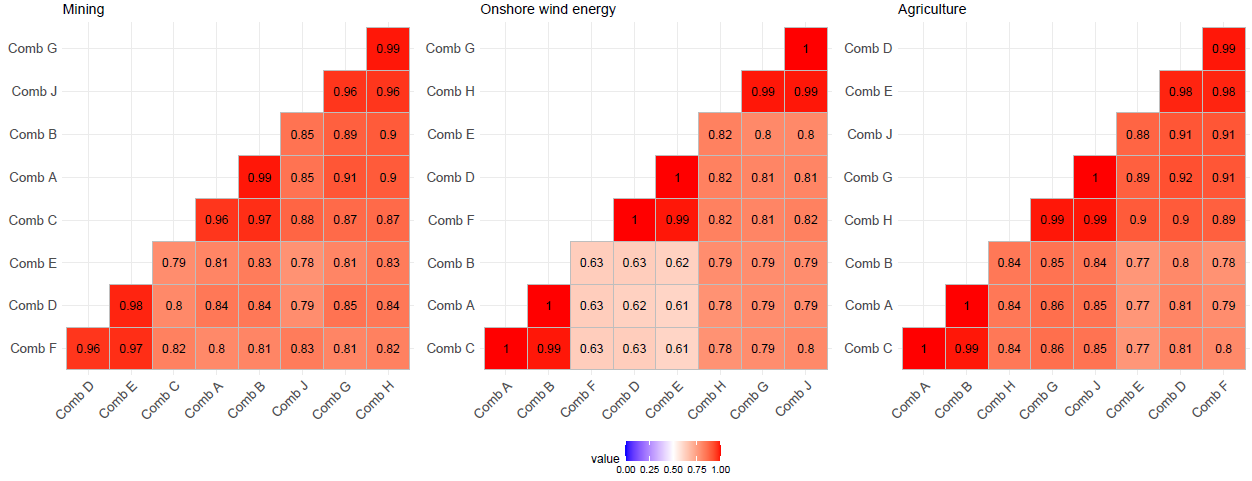

Supplementary Figure 1. Spearman rank correlations between metric combinations A-H with respect to ranking sites for ecological sensitivity for Mining, Onshore wind energy and Agriculture. (Sensitivity thresholds were based on the upper quartile definition and metric scores combined using the ‘Trigger’ scoring system.)

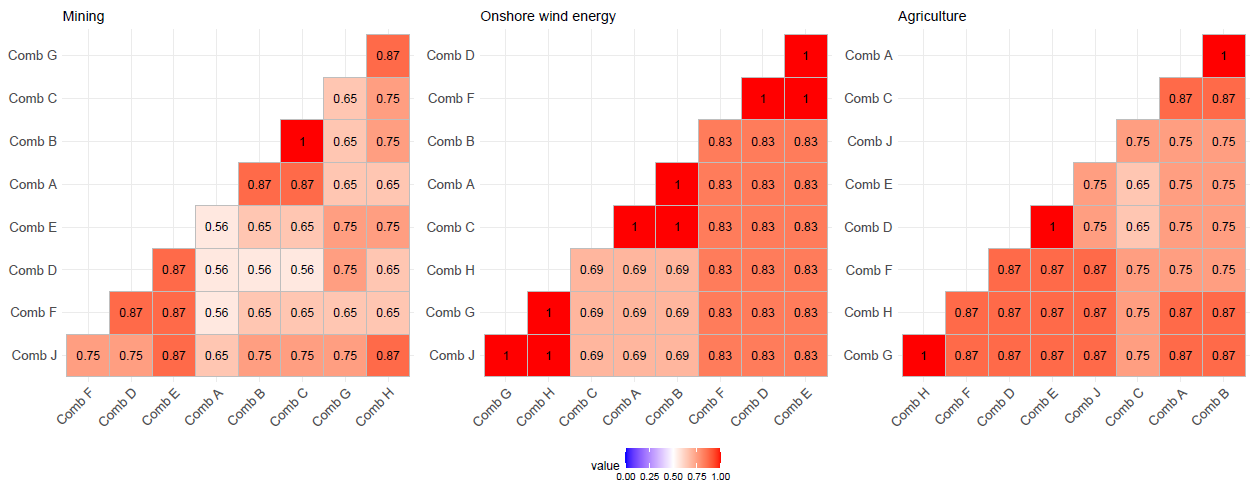

Supplementary Figure 2. Jaccard similarity matrices for metric combinations A-H with respects to sites ranked in the top 20% for ecological sensitivity for Mining, Onshore wind energy and Agriculture. (Sensitivity thresholds were based on the upper quartile definition and metric scores combined using the ‘Trigger’ scoring system.)

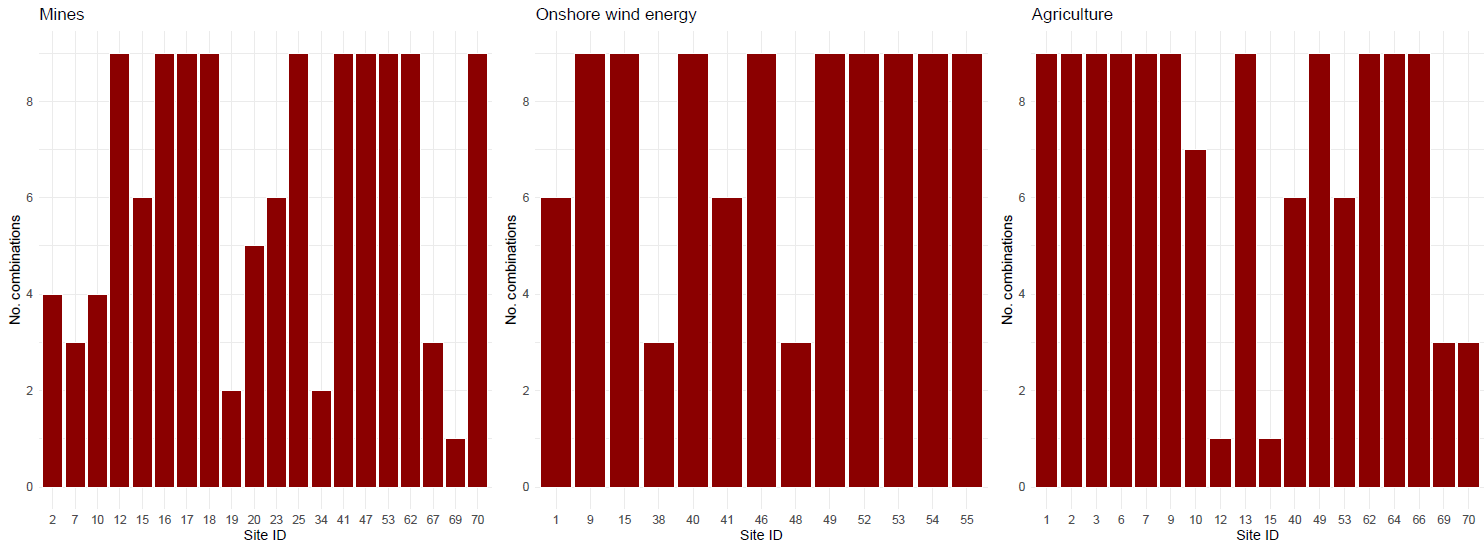

Supplementary Figure 3. Frequency chart showing the number of metric combinations from combinations A-J for which an individual site was ranked in the top 20% with respect to ecological sensitivity for Mining, Onshore wind energy and Agriculture. (Sensitivity thresholds were based on the upper quartile definition and metric scores combined using the ‘Trigger’ scoring system.)

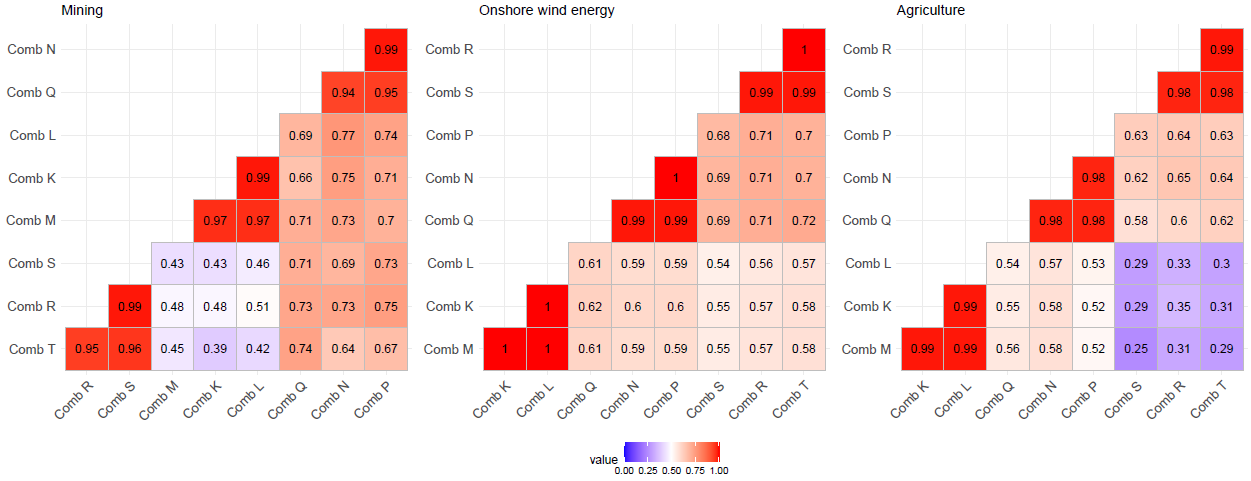

Supplementary Figure 4. Spearman rank correlations between metric combinations K-T with respect to ranking sites for ecological sensitivity including restoration potential for Mining, Onshore wind energy and Agriculture. (Sensitivity thresholds were based on the upper quartile definition and metric scores combined using the ‘Trigger’ scoring system.)

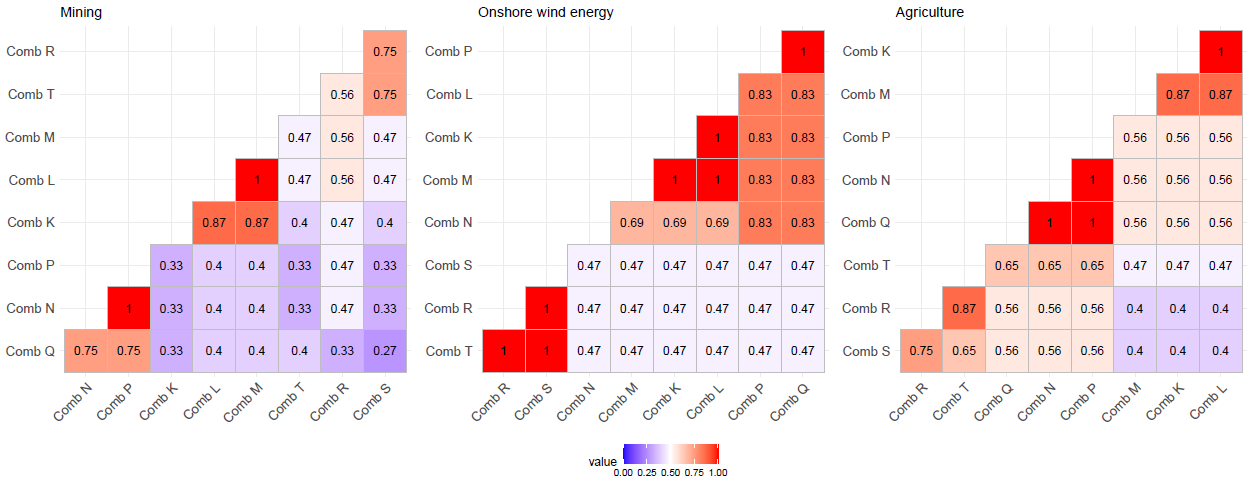

Supplementary Figure 5. Jaccard similarity matrices for metric combinations K-T with respect to sites ranked in the top 20% for ecological sensitivity including restoration potential for Mining, Onshore wind and Agriculture. (Sensitivity thresholds were based on the upper quartile definition and metric scores combined using the ‘Trigger’ scoring system.)

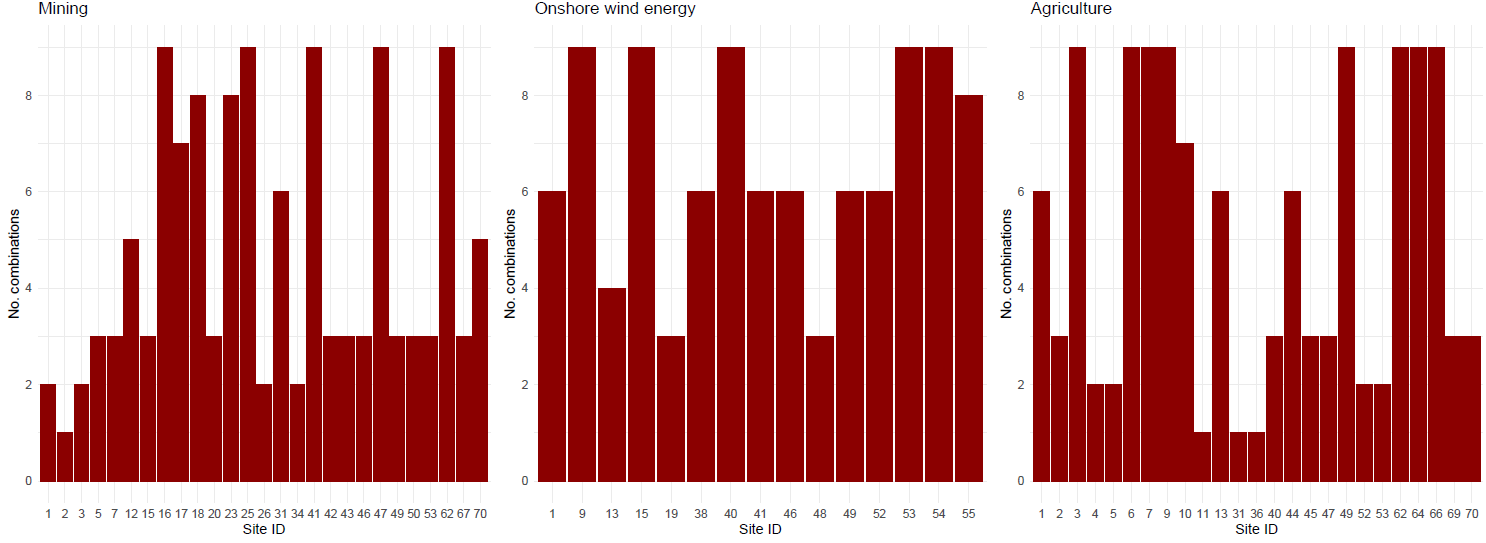

Supplementary Figure 6. Frequency chart showing the number of metric combinations from combinations K-T for which an individual site was ranked in the top 20% with respect to ecological sensitivity including restoration potential for Mining, Onshore wind energy and Agriculture. (Sensitivity thresholds were based on the upper quartile definition and metric scores combined using the ‘Trigger’ scoring system.)

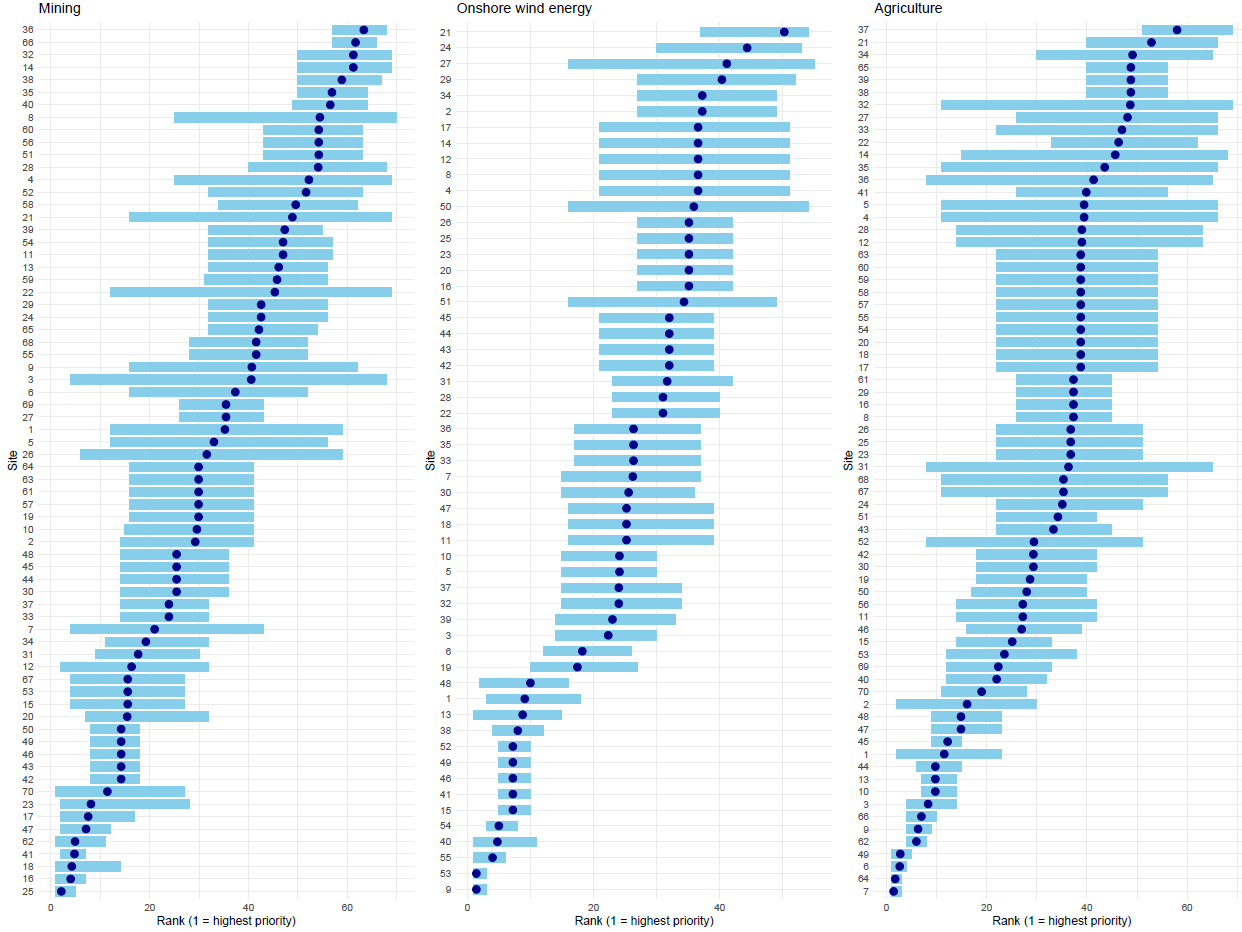

Supplementary Figure 7. Rank shift plots showing the mean ecological sensitivity and restoration potential ranking (dark blue dot) and maximum and minimum ranking (blue line) for each site based on metric combinations K-T for Mining, Onshore wind energy and Agriculture. (Sensitivity thresholds were based on the upper quartile definition and metric scores combined using the ‘Trigger’ scoring system.)

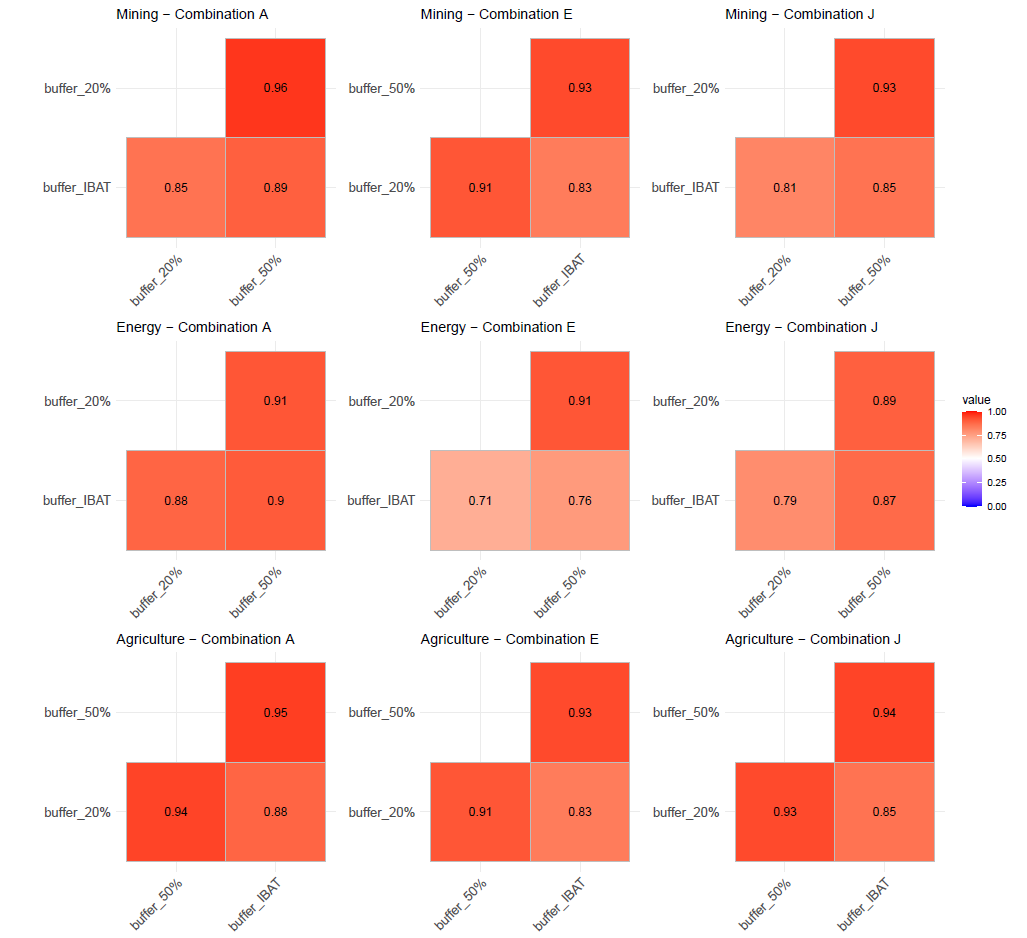

Supplementary Figure 8. Spearman rank correlations between buffer sizes for metric combinations A, E and J with respect to ranking sites for ecological sensitivity for Mining, Onshore wind energy and Agriculture. (Sensitivity thresholds were based on IBAT recommendations (proximity to PAs and KBAs, STAR_t_) or the upper quartile definition (all other metrics) and metrics were combined using the Trigger scoring system.)

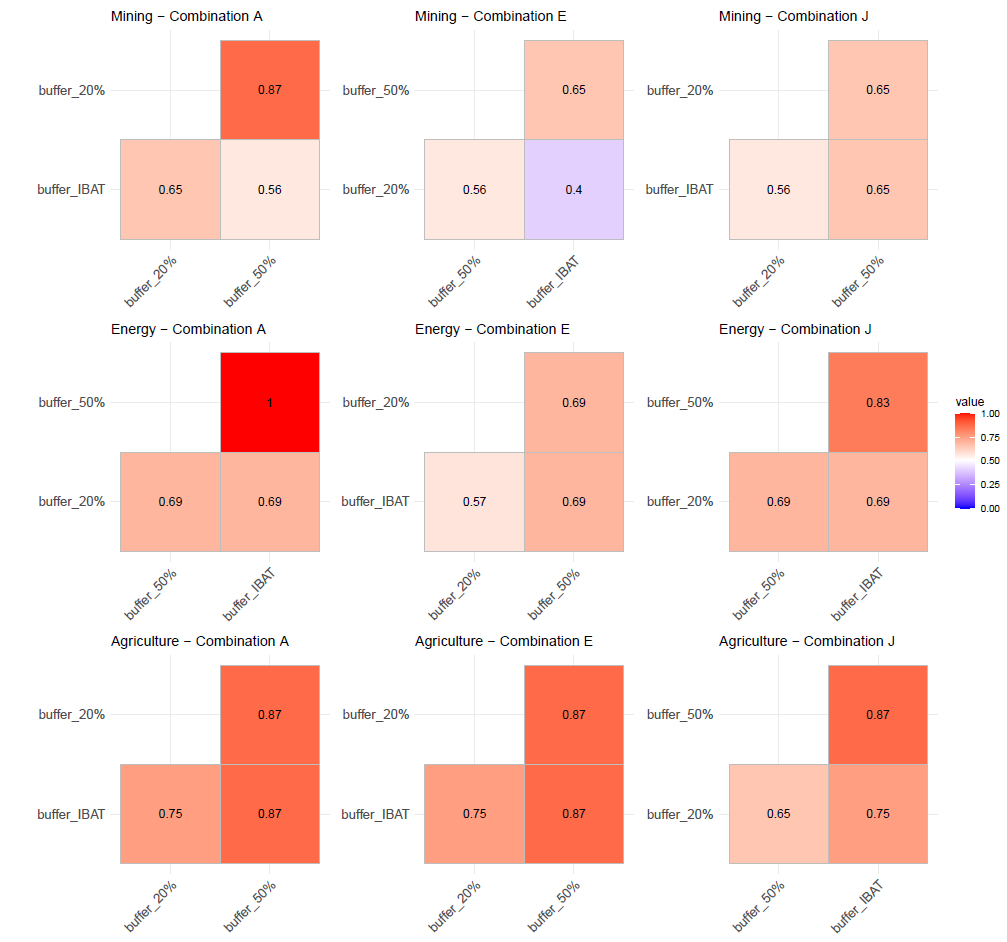

Supplementary Figure 9. Jaccard similarity matrices between buffer sizes with respect to sites ranked in the top 20% for ecological sensitivity by metric combinations A, E and J for Mining, Onshore wind energy and Agriculture. (Sensitivity thresholds were based on IBAT recommendations (proximity to PAs and KBAs, STAR_t_) or the upper quartile definition (all other metrics) and metrics were combined using the Trigger scoring system.)

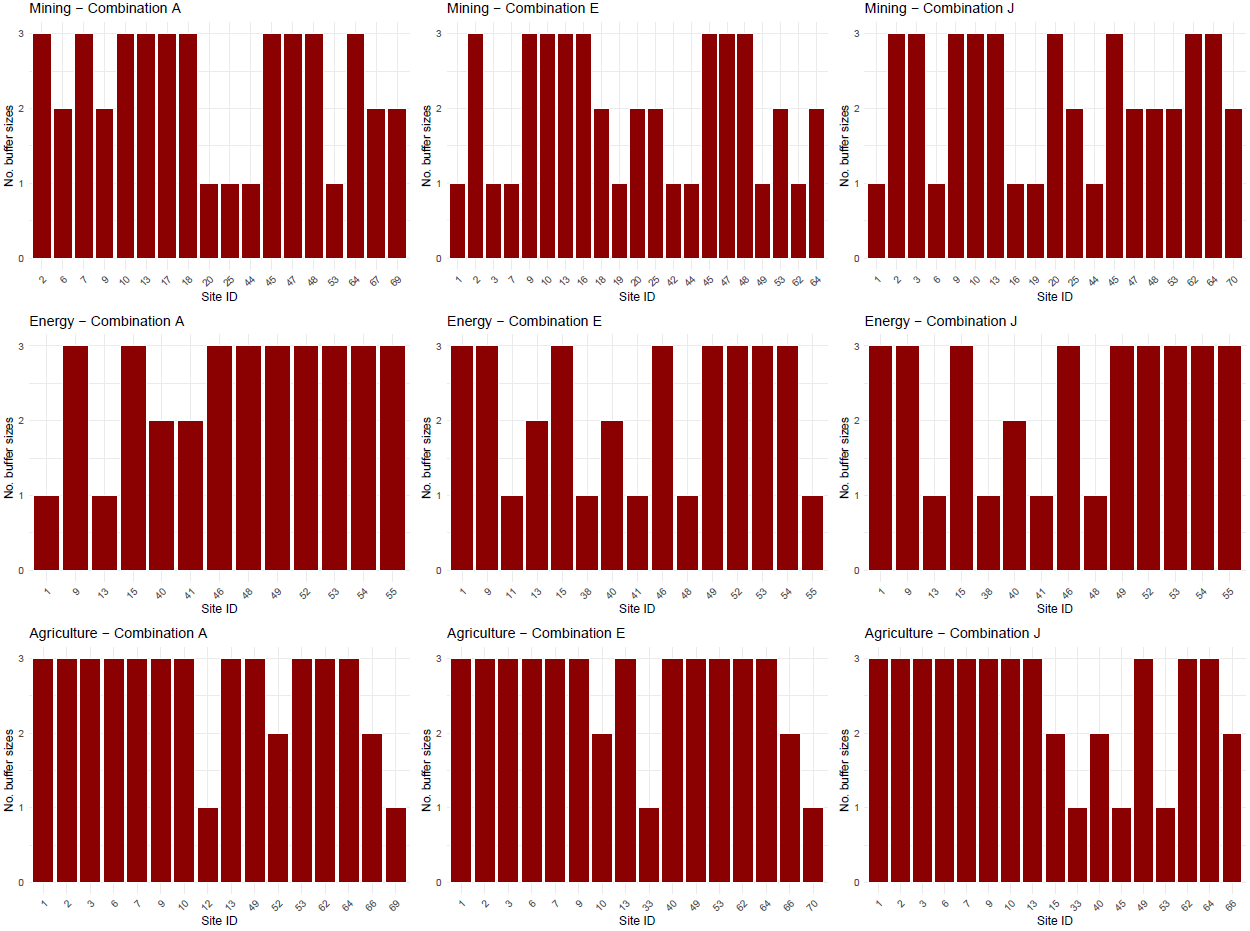

Supplementary Figure 10. Frequency chart showing the number of buffer sizes which ranked an individual site in the top 20% with respect to ecological sensitivity for Mining, Onshore wind energy and Agriculture using metric combinations A, E and J. (Sensitivity thresholds were based on IBAT recommendations (proximity to PAs and KBAs, STAR_t_) or the upper quartile definition (all other metrics) and metrics were combined using the Trigger scoring system.)

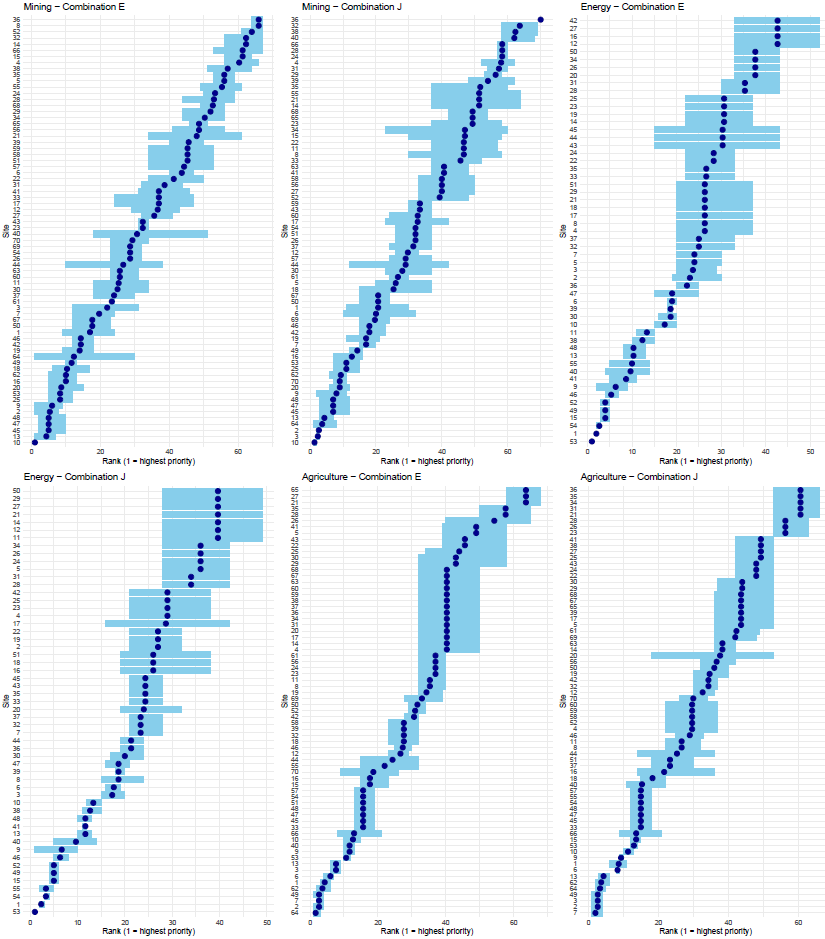

Supplementary Figure 11. Rank shift plots showing the mean ecological sensitivity ranking (dark blue dot) and maximum and minimum ranking (blue line) for each site for the three buffer sizes for metric combinations E and J for mining, onshore wind and agriculture. (Sensitivity thresholds were based on IBAT recommendations (proximity to PAs and KBAs, STAR_T_) or the upper quartile definition (all other metrics) and metrics were combined using the Trigger scoring system.)

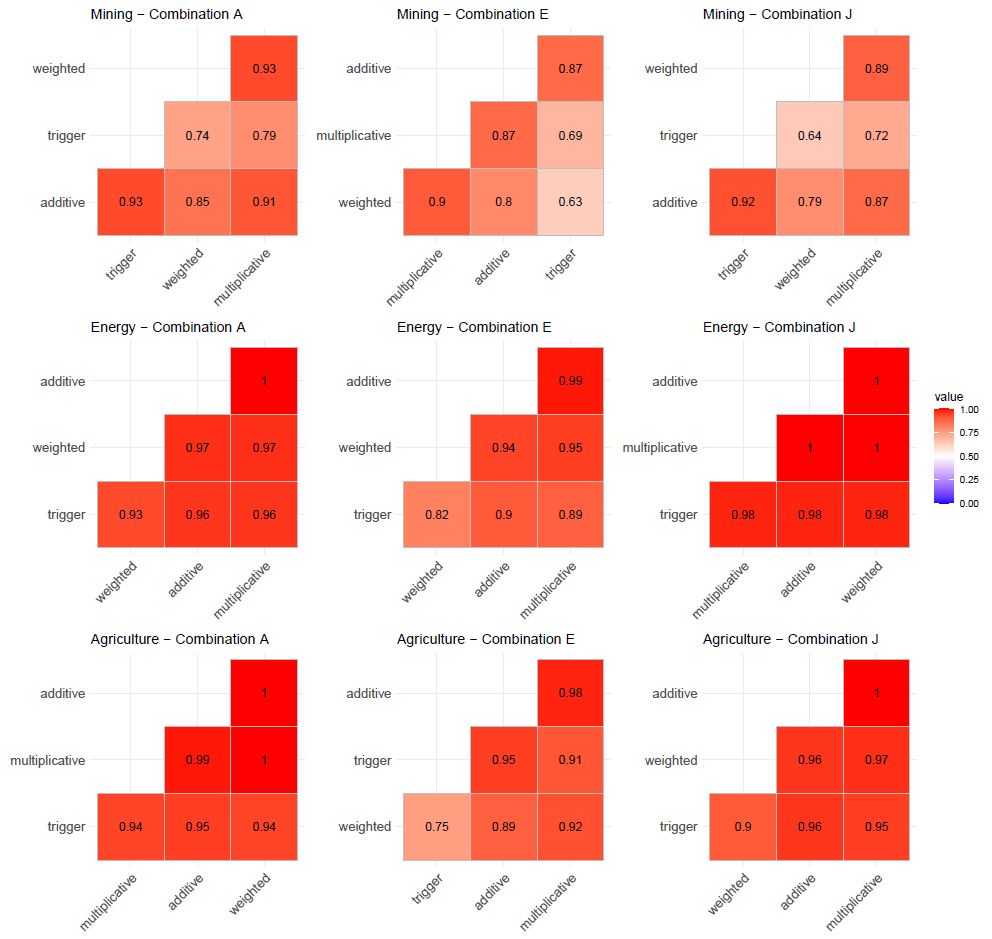

Supplementary Figure 12. Spearman rank correlations between scoring systems for metric combinations A, E and J with respect to ranking sites for ecological sensitivity for Mining, Onshore wind energy and Agriculture. (Sensitivity thresholds were based on the upper quartile definition.)

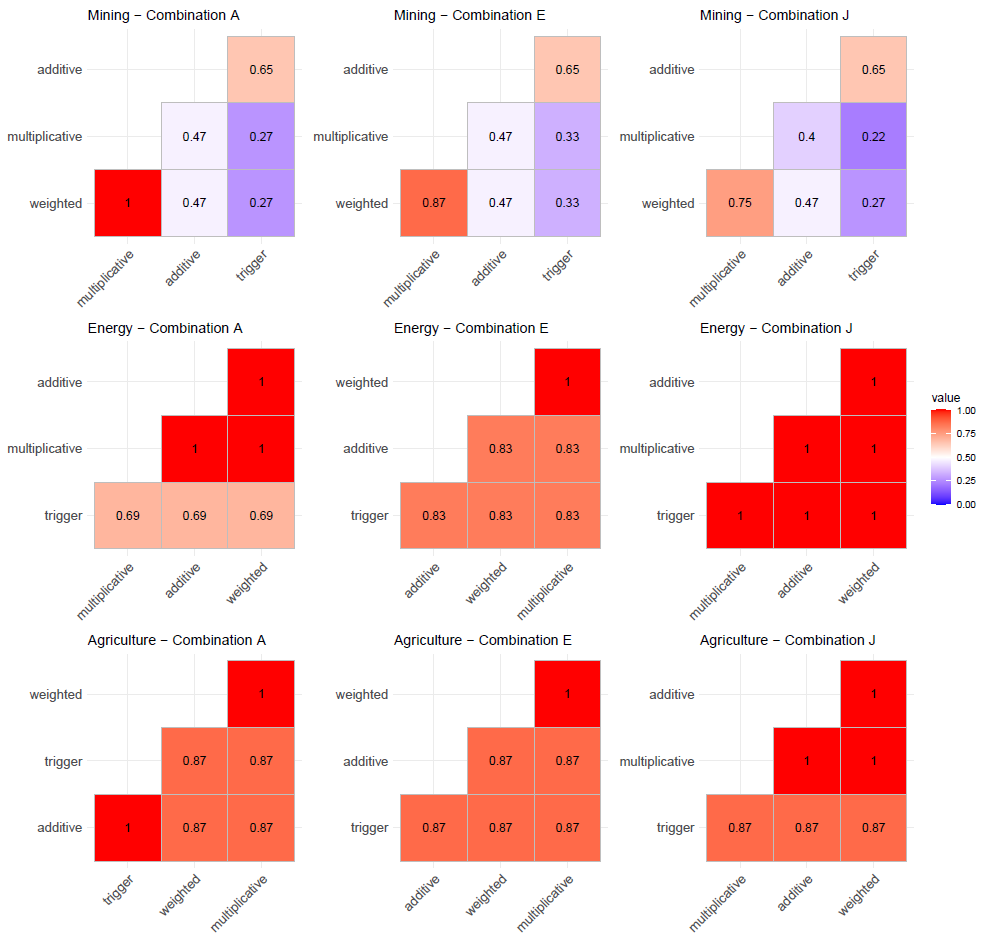

Supplementary Figure 13. Jaccard similarity matrices between scoring systems with respect to sites ranked in the top 20% for ecological sensitivity by metric combinations A, E and J for Mining, Onshore wind energy and Agriculture. (Sensitivity thresholds were based on the upper quartile definition.)

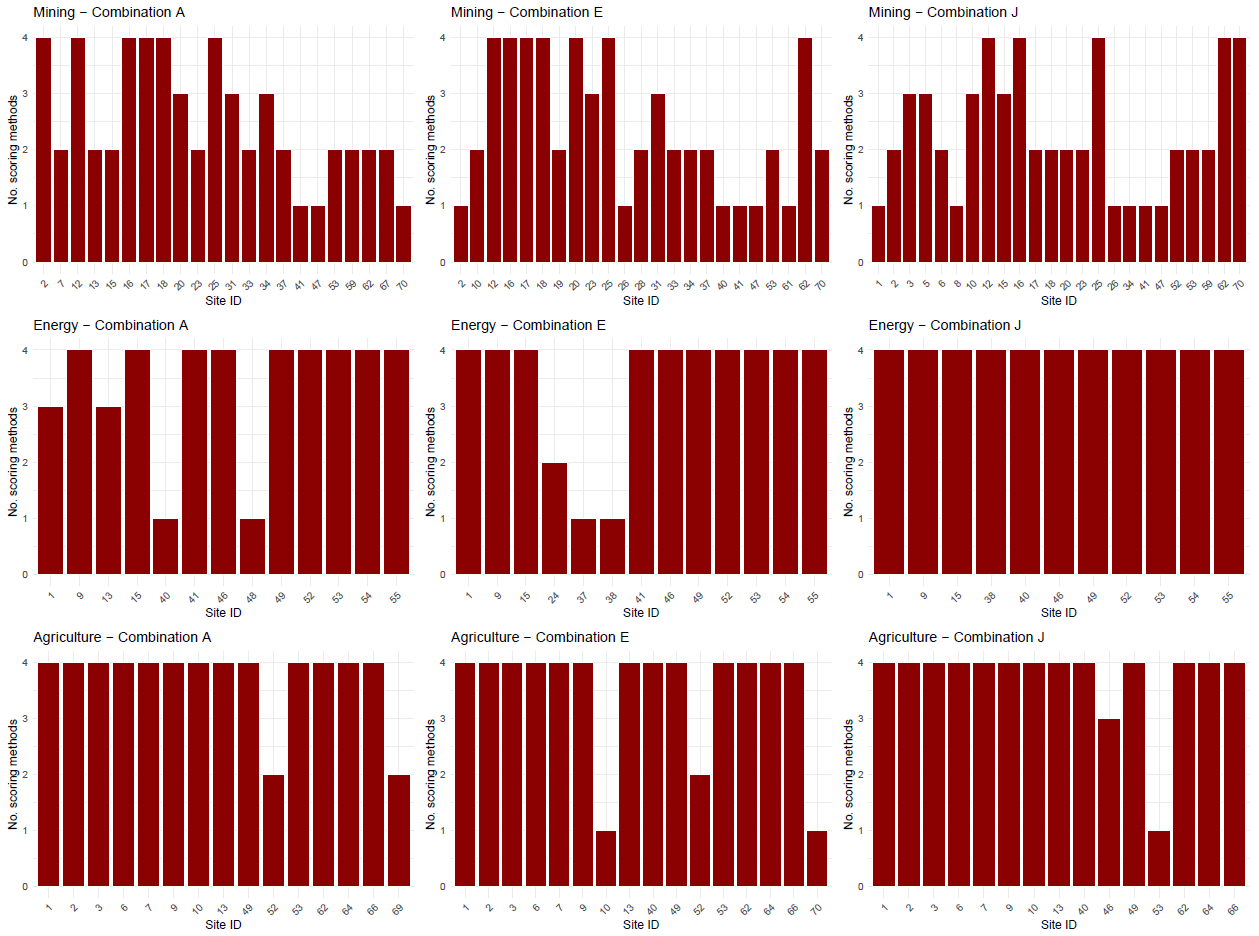

Supplementary Figure 14. Frequency chart showing the number of scoring methods which ranked an individual site in the top 20% with respect to ecological sensitivity for Mining, Onshore wind and Agriculture using metric combinations A, E and J. (Sensitivity thresholds were based on the upper quartile definition.)

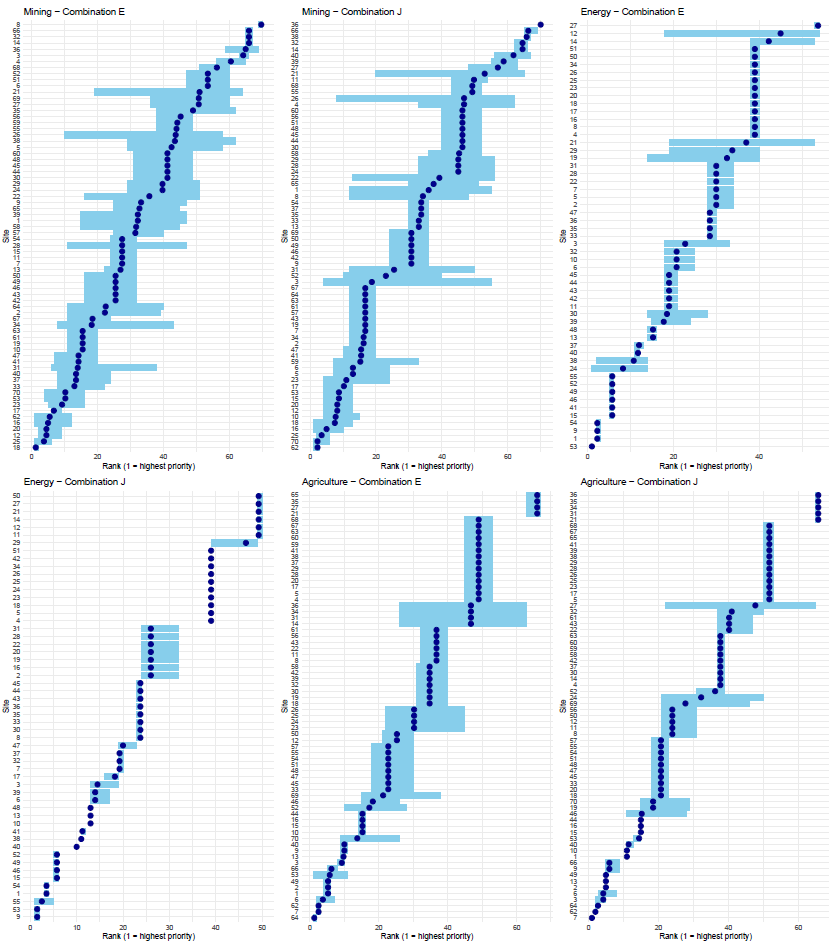

Supplementary Figure 15. Rank shift plots showing the mean ecological sensitivity ranking (dark blue dot) and maximum and minimum ranking (blue line) for each site for the four scoring systems for metric combinations E and J for Mining, Onshore wind energy and Agriculture. (Sensitivity thresholds were based on the upper quartile definition.)

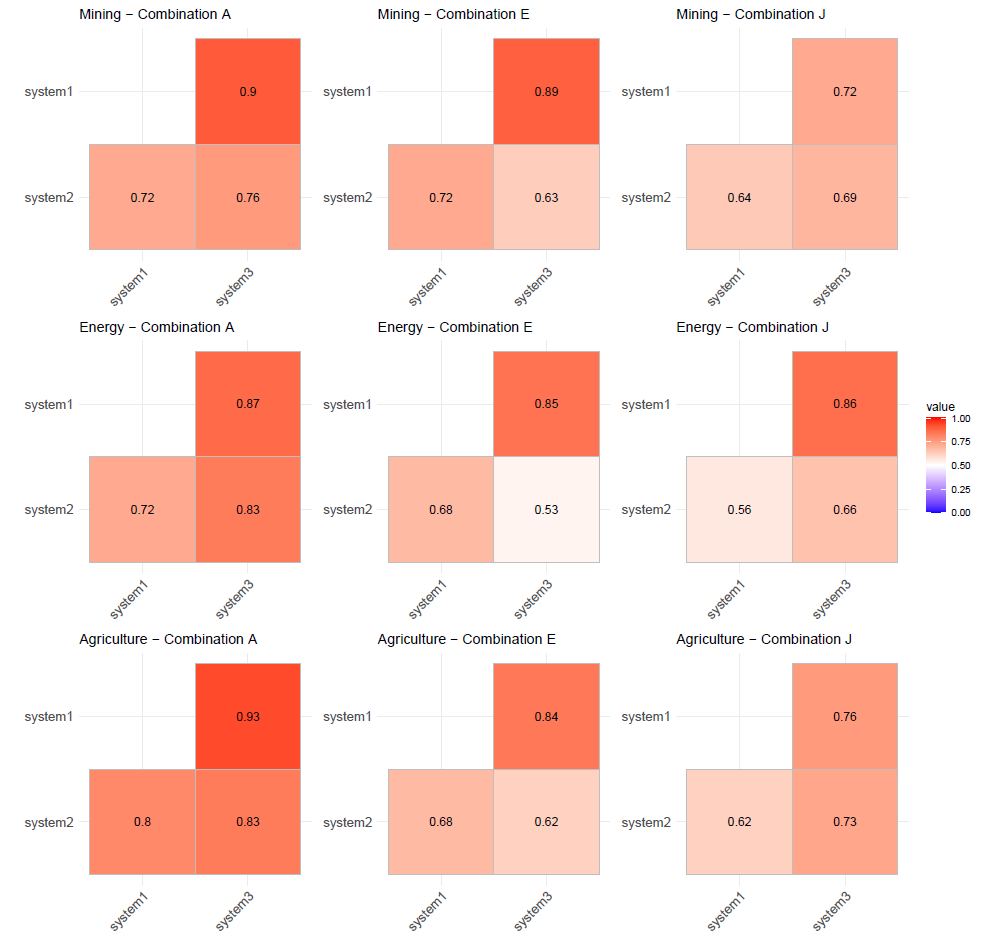

Supplementary Figure 16. Spearman rank correlations between sensitivity threshold definitions for metric combinations A, E and J with respect to ranking sites for ecological sensitivity for Mining, Onshore wind energy and Agriculture. (Metrics were combined using the trigger scoring system.)

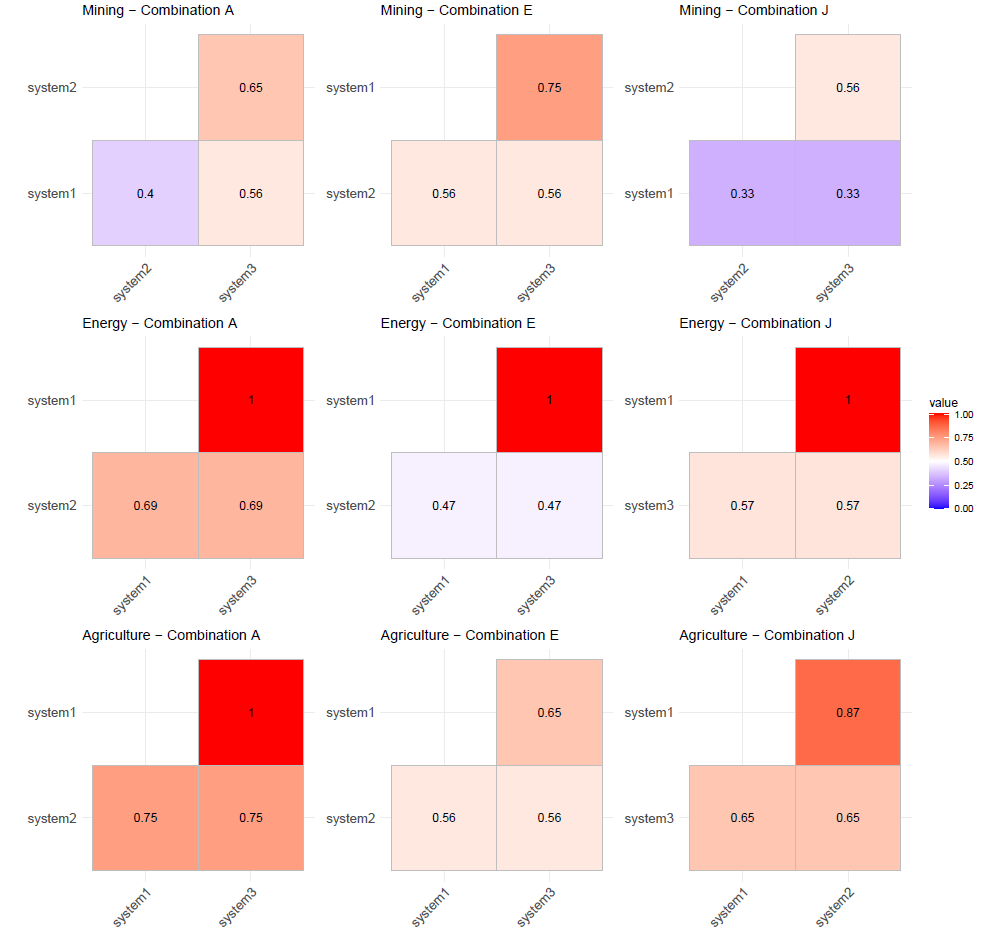

Supplementary Figure 17. Jaccard similarity matrices between sensitivity threshold definitions with respect to sites ranked in the top 20% for ecological sensitivity by metric combinations A, E and J for Mining, Onshore wind energy and Agriculture. (Metrics were combined using the Trigger scoring system.)

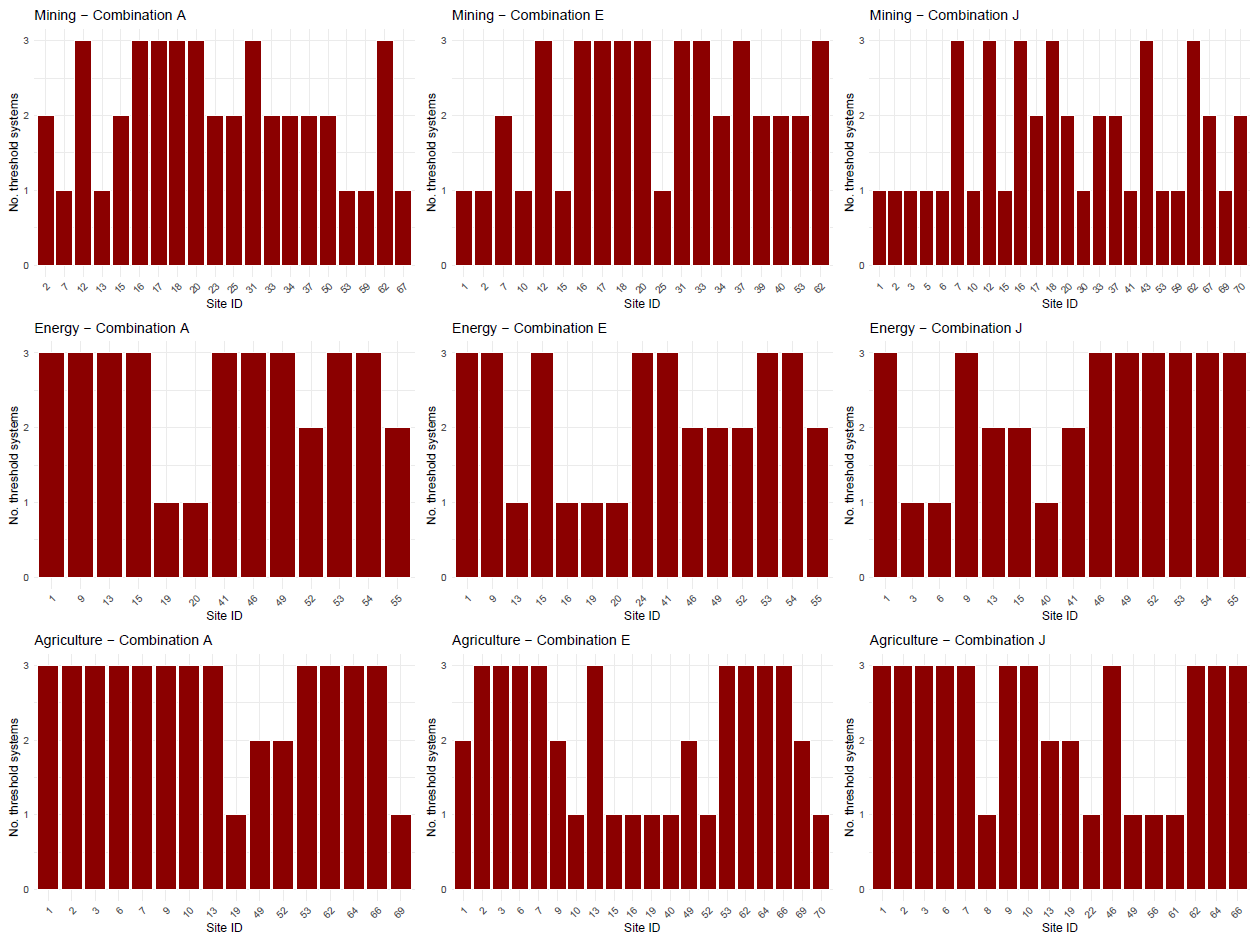

Supplementary Figure 18. Frequency chart showing the number sensitivity threshold definitions which ranked an individual site in the top 20% with respect to ecological sensitivity for Mining, Onshore wind energy and Agriculture using metric combinations A, E and J. (Metrics were combined using the Trigger scoring system.)

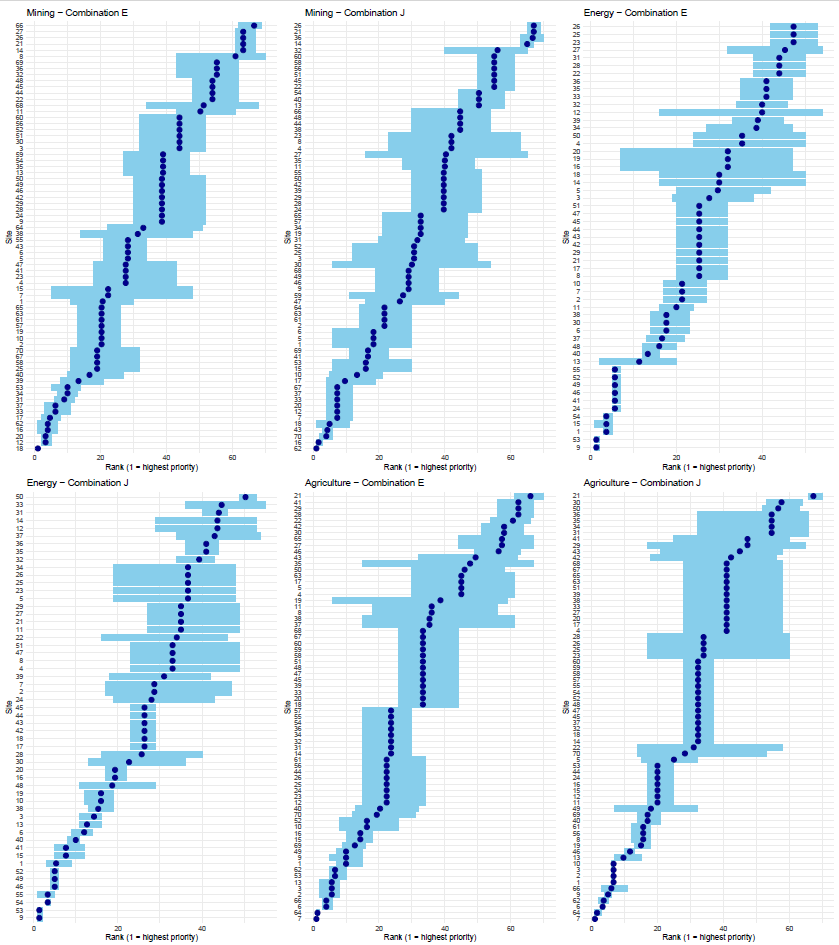

Supplementary Figure 19. Rank shift plots showing the mean ecological sensitivity ranking (dark blue dot) and maximum and minimum ranking (blue line) for each site for the four sensitivity threshold definitions for metric combinations E and J for Mining, Onshore wind energy and Agriculture. (Metrics were combined using the trigger scoring system.)
